# E3 ubiquitin ligase RNF213 employs a non-canonical zinc finger active site and is allosterically regulated by ATP

**DOI:** 10.1101/2021.05.10.443411

**Authors:** Juraj Ahel, Adam Fletcher, Daniel B. Grabarczyk, Elisabeth Roitinger, Luiza Deszcz, Anita Lehner, Satpal Virdee, Tim Clausen

**Affiliations:** Research Institute of Molecular Pathology (IMP), Vienna BioCenter, Vienna, Austria; MRC Protein Phosphorylation and Ubiquitylation Unit, School of Life Sciences, University of Dundee, Dundee, United Kingdom; Institute of Molecular Biotechnology (IMBA), Vienna BioCenter, Vienna, Austria; Vienna BioCenter Core Facilities, Vienna BioCenter, Vienna, Austria.; Medical University of Vienna, Vienna, Austria

## Abstract

RNF213 is a giant E3 ubiquitin ligase and a major susceptibility factor of Moyamoya disease, a cerebrovascular disorder that can result in stroke or death. In the cell, RNF213 is involved in lipid droplet formation, lipotoxicity, hypoxia, and NF-κB signaling, but its exact function in these processes is unclear. Structural characterization has revealed the presence of a dynein-like ATPase module and an unprecedented but poorly understood E3 module. Here, we demonstrate that RNF213 E3 activity is dependent on ATP binding, rather than ATP hydrolysis, and is particularly responsive to the ATP/ADP/AMP ratio. Biochemical and activity-based probe analyses identify a non-canonical zinc finger domain as the E3 active site, which utilizes the strictly conserved Cys4462, not involved in zinc coordination, as the reactive nucleophile. The cryo-EM structure of the trapped RNF213:E2∼Ub intermediate reveals RNF213 C-terminal domain as the E2 docking site, which positions the ubiquitin-loaded E2 proximal to the catalytic zinc finger, facilitating nucleophilic attack of Cys4462 on the E2∼Ub thioester. Our findings show that RNF213 represents an undescribed type of a transthiolation E3 enzyme and is regulated by adenine nucleotide concentration via its ATPase core, possibly allowing it to react to changing metabolic conditions in the cell.

## Introduction

RNF213/ALO17/mysterin (Ring Finger Protein 213) is the largest known human single-chain E3 ubiquitin ligase, first described as the major susceptibility factor for Moyamoya disease (MMD) (*1*), a disorder which causes aberrant vasculature development commonly leading to stroke and other complications (*2*). Apart from MMD, RNF213 has been implicated in other vascular pathologies like intracranial atherosclerosis (*3*) and in angiogenesis, as shown by studies in mice (*4*), zebrafish (*5*), and tissue culture (*4, 6*). Connections to non-vascular diseases were also reported, including diabetes (*7*), cancer (*8*) and both viral (*9*) and bacterial (*10*) infection. However, the molecular basis for this wide range of pathologies remains unknown. In the cell, RNF213 was shown to be involved in fatty acid metabolism (*11*), and to associate with intracellular lipid droplets in a manner dependent on its E3 ligase and ATPase activities (*12*). Moreover, RNF213 was demonstrated to regulate non-mitochondrial oxygen consumption (*13*), consequently playing a role in tumor resistance to hypoxia, and to be capable of activating the NF-κB signaling pathway (*11, 13, 14*), hinting at involvement in the immune response. The *RNF213* gene is also the gene with the most strongly accelerated rate of evolution along the ancestral branch leading to hominines, suggestive of a key role in large brain development (*15*).

Despite these insights, the molecular mechanism of RNF213 is only starting to be understood. We recently described the structure of *M. musculus* RNF213 (PDB: 6TAX) and characterized its enzymatic activity, revealing it as a large monomeric protein with a functional dynein-like AAA (ATPases Associated with diverse cellular Activities) core that exists alongside a novel E3 ligase module (*16*). However, despite its demonstrated E3 activity, the catalytic mechanism of this E3 module remains unclear. Canonically, E3 ubiquitin ligases are classified as one of the three major types: RING (Really Interesting New Gene), HECT (Homologous to E6-AP Carboxyl Terminus), or RBR (RING-between-RING). RING E3s contain a zinc-finger domain that coordinates two metal ions with cross-brace architecture eponymously termed the RING domain (*17*). Members of this family recruit substrates via their non-RING domains and act as catalytic scaffolds whereby they facilitate Ub transfer from the upstream thioester-linked E2- ubiquitin intermediate (E2∼Ub) to the substrate. In contrast, HECT and RBR E3 ligases first accept Ub from E2∼Ub onto an acceptor cysteine - a process termed transthiolation - from where it is typically transferred to a lysine side chain by aminolysis reaction, thereby forming an isopeptide bond (*18, 19*). Divergence from these well-established E3 mechanisms was evidenced with the large E3 ligase MYCBP2, which employs a RCR (RING-Cys-Relay) mechanism involving two acceptor cysteines (*20*). The first cysteine undergoes transthiolation with E2∼Ub and Ub is then relayed to the second cysteine. The active site of the second cysteine confers esterification activity towards model substrates, with high selectivity for threonine over serine. This highlights the unexplored mechanistic diversity and substrate scope of the ubiquitin system and suggests that additional E3 subtypes are likely to emerge.

A number of publications reported that the isolated RING domain from RNF213 has E3 activity (*1*) that works with either the E2 UBE2D2/UbcH5b (*21*), or the E2 UBE2N/Ubc13 (*14, 22*). However, our recent report (*16*) reveals that the RING domain is not required for basal E3 auto- ubiquitination activity of full-length RNF213. As RING is the only domain within RNF213 associated with E3 activity, an uncharacterized element conferring the auto-ubiquitination activity must exist. Furthermore, our observed auto-ubiquitination activity is most efficient with the E2 UBE2L3/UbcH7, which cannot support RING E3 activity because it lacks features required for lysine positioning and/or deprotonation (*18, 23*). These findings indicate that RNF213 might contain a catalytic cysteine nucleophile (reminiscent of HECT, RBR, and RCR subtypes) that undergoes transthiolation with E2∼Ub. Nevertheless, further studies are required to elucidate the functional relevance of the two distinct E3-associated domains within RNF213. Likewise, it is an open question whether other domain(s) of RNF213 are involved in E3 ligase activity and how the ubiquitination activity might be coupled with the ATPase function of RNF213. Our previous cryo-EM analysis illustrated the structural organization of the E3 and ATPase modules, which are complemented by an N-terminal stalk region of unknown function (**Figure 2a**), and provided a structural framework for understanding the interplay between all the subdomains of this protein. In the present study, we continued this analysis and identified the E3 active site – including the nucleophile mediating the transthiolation reaction – and established that the E3 transthiolation activity is regulated by the attached ATPase core.

**Figure 1.**
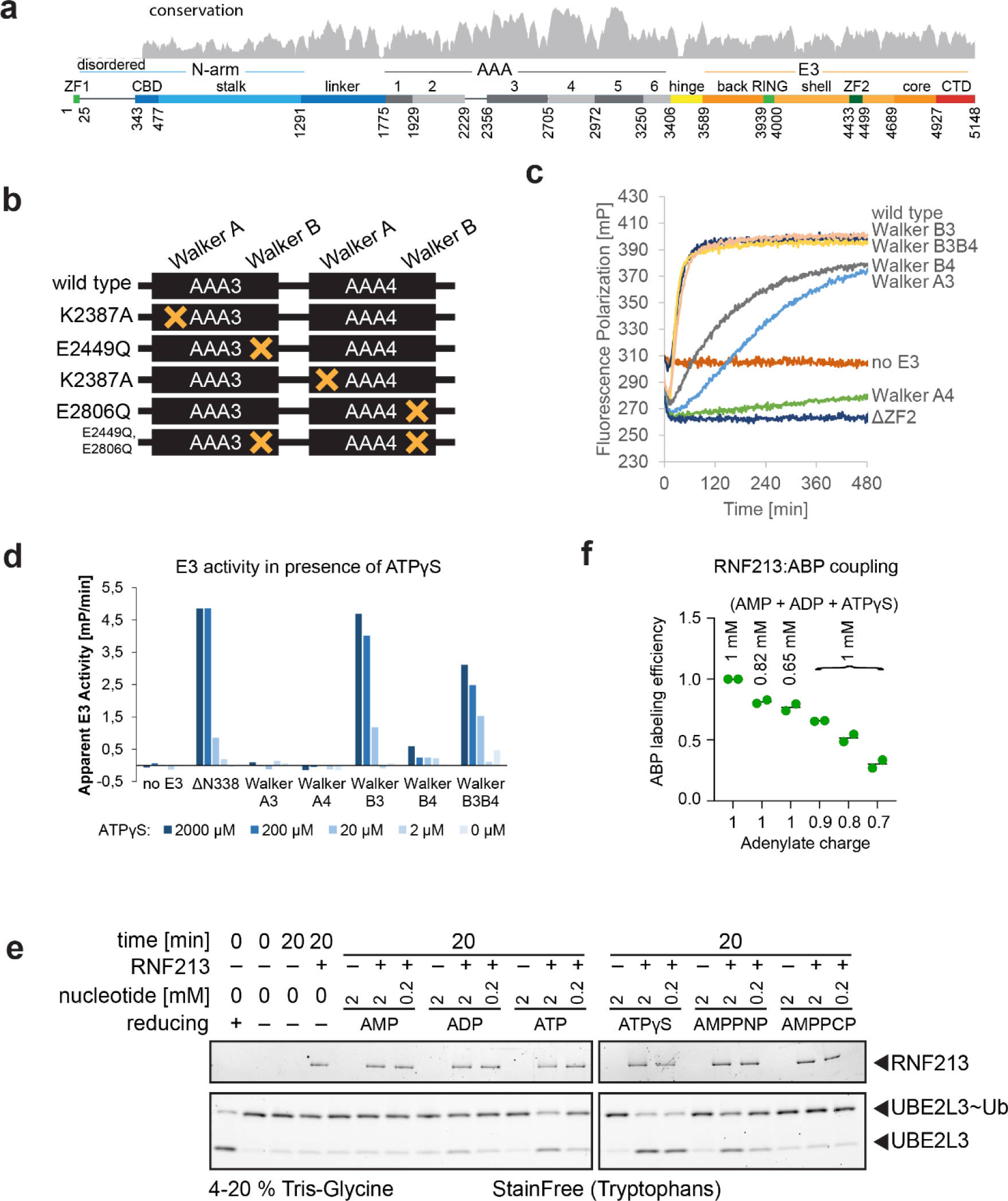
Nucleotide regulation of the E3 activity of RNF213. (a) A schematic overview of RNF213, indicating the known and predicted functional domains. The numbers below the schematic indicate start and end residues of the indicated domain or motif. Evolutionary conservation profile is shown above. (b) Illustration of the analyzed ATPase mutants of RNF213. (c) Example timecourse curves for the FP-based E3 activity assay. Reactions are carried out with RNF213 in presence of ubiquitin partially labeled with DyLight488. (d) Activity of ATPase mutants compared to the wild type (ΔN338) protein. Walker A mutations lead to near-full loss of activity. Walker B mutations do not prominently affect activity. Change in polarization over time is used as the measure of apparent E3 activity. (e) UBE2L3∼Ub discharge assay in presence of various nucleotides. When ATP analogues are present, depletion of the UBE2L3∼Ub band and appearance of the free UBE2L3 band can be observed, except when the analogue is AMPPCP. With ADP, only a very slight increase in lower band intensity is observable, while AMP alone has no detectable effect on discharge rate. Example timecourses are given in **Figure1-figure supplement 1**. Additional quantifications of E3 activity in presence of different nucleotides, raw data for RNF213-ABP coupling, dependence of E3 activity on the concentration of ATPγS, and a competition assay with ATPgS and AMP or ADP are given in **Figure1-figure supplement 2**. E3 activity in presence of nucleoside triphosphates of bases other than adenine is presented in **Figure1-figure supplement 3.** (f) Qualitative analysis of RNF213-ABP coupling efficiency at various levels of adenylate charge, using ATPγS as a substitute for ATP. The first three reactions contain equivalent amounts of ATPγS as the last three, but without added ADP or AMP. Prominent reduction of activity is only present upon adding the ADP/AMP, and not when ATPγS concentration is simply lowered. Two independent measurements for each condition are displayed.

**Figure 2.**
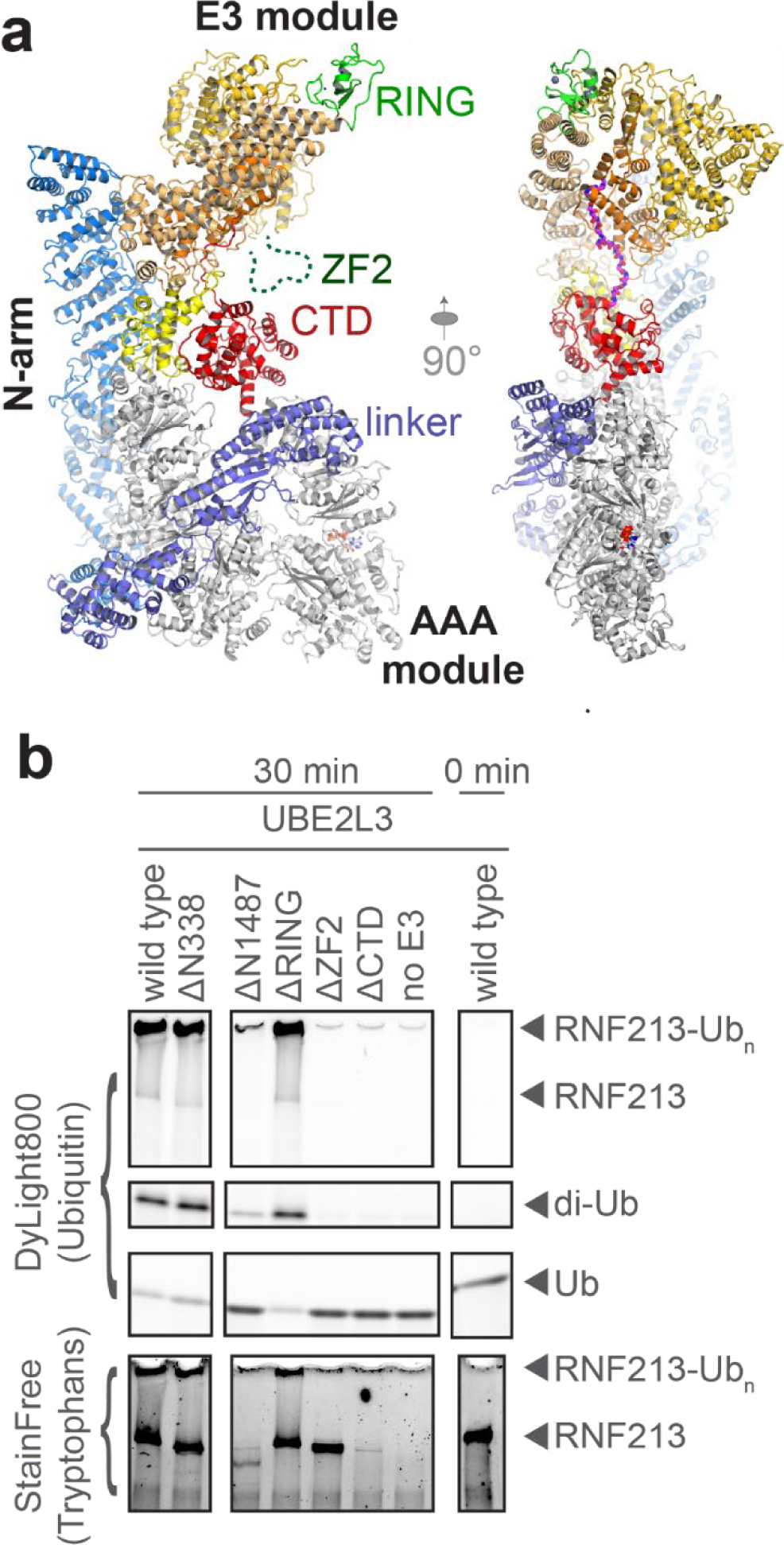
Role of different domains of RNF213 in its E3 ligase activity. (a) Overview of the 3D structure of RNF213, shown as a ribbon model. Side view (left) and front view (right). Coloring of the domains corresponds to the coloring in the sequence overview in Figure 1a. (b) E3 ligase assay with RNF213 domain deletion mutants. The N-terminal stalk appears entirely dispensable for auto-ubiquitination, as does the RING domain. Deletion of ZF2 or CTD leads to near-complete loss of activity with UBE2L3. Concentration of RNF213 in reactions containing the ΔN1487 and ΔCTD variants is lower than in other reactions, but still permits qualitative assessment of activity. Figure 2-figure supplement 1 shows the activity of the variants in presence of UBE2D3 or in absence of an E2.

### Regulation of RNF213 activity

RNF213 is a large multi-domain protein, with a dynein-like AAA core (**Figure 1a**). Mutations of the ATPase active sites have been reported to affect RNF213 oligomerization (*5*), and the ability of RNF213 to associate with intracellular lipid droplets (*12*). Despite this, it remains unclear how this regulation of behavior occurs, or what biological role it serves.

To address whether the AAA ATPase activity is coupled with the E3 activity, we mutated the two functional AAA3 and AAA4 ATPases in the dynein-like core, disrupting either ATP binding (Lys>Ala mutation in the Walker A motif) or ATP hydrolysis (Glu>Gln mutation in the Walker B motif) (*24*) (**Figure 1b**). Remarkably, Walker A mutants are almost entirely inactive in auto-ubiquitination assays, contrasting the Walker B mutants which retain high E3 ligase activity (**Figure 1c,**d). This shows that ATP binding to AAA3 and AAA4 is both necessary and sufficient to promote the E3 ligase activity of RNF213, while ATP hydrolysis at those sites has no major effect on activity, at least when the E3 substrate is RNF213 itself.

If ATP binding to RNF213 is necessary and sufficient to induce RNF213 activity, then wild-type RNF213 should be inactive in the absence of ATP, but get activated in the presence of a non-hydrolyzable ATP analog. When ATP is substituted with the poorly-hydrolyzable ATPγS – a nucleotide analog that can be also used by the E1 enzyme in the ubiquitination assay (*25*) - RNF213 indeed not only remains active, but has an increased auto-ubiquitination rate, confirming that ATP binding rather than hydrolysis stimulates its E3 activity (**Figure 1e**). We tested this mode of regulation further in an E2-Ub discharge assay which provides more defined reaction conditions by omitting the E1 enzyme. For this purpose, we purified the thioester- linked UBE3L3∼Ub intermediate, mixed it with RNF213 and various adenosine nucleotides (AMP, ADP, ATP, ATPγS, AMP-PNP, AMP-PCP) and assayed the discharge of Ub from the E2 enzyme. RNF213 shows no enhancement of UBE2L3∼Ub discharge in the absence of a nucleotide, demonstrating that ATP binding is strictly required for ubiquitin transfer (**Figure 1-figure supplement 1**). Furthermore, non-hydrolyzable analogs ATPγS and AMP-PNP are also able to stimulate RNF213 activity, even more strongly than ATP itself (**Figure 1e**, **Figure 1-figure supplement 2b**). ADP stimulates the activity very weakly, while with AMP-PCP or AMP RNF213 remains inactive (**Figure 1e**). We also tested other nucleoside triphosphates and found that only adenosine-based nucleosides efficiently stimulate RNF213 activity (**Figure 1- figure supplement 3**), solidifying the claim that RNF213 is specifically regulated by ATP.

UBE2L3 has been described to be only reactive towards cysteine residues (*18*), so we hypothesized that RNF213 employs a ubiquitin transfer mechanism similar to the mechanism of HECT, RBR, or RCR E3 ligases. Hinting at RNF213 utilizing a catalytic cysteine-dependent transthiolation mechanism, RNF213 was detected in UBE2L3 activity-based proteomic profiling of a human neuroblastoma cell lysate (*20*). To further explore this possibility, we labeled RNF213 with an activity-based probe (ABP) based on the upstream UBE2L3∼Ub conjugate, which can form a stable covalent product with the Ub acceptor nucleophile of cysteine-mediated E3 ligases (*26*) (**Figure 1-figure supplement 4**). Consistent with this, a UBE2L3 activity-based probe efficiently labelled RNF213 in a gel-based assay only in the presence of a suitable nucleotide (**Figure 1-figure supplement 1a**). Notably, as the ABP only assesses E2-E3 transthiolation activity, this demonstrates that the nucleotide-mediated RNF213 regulation operates at the level of E2-E3 transthiolation.

To further clarify the dependence of RNF213 E3 activity on ATP binding, we performed competition assays with ATPγS and ADP or AMP. Increasing concentrations of ADP and AMP led to lower activity of RNF213, indicating that both species are inhibitory, but with a stronger effect for AMP. This might be mediated by competitive inhibition of the same AAA active sites that bind ATPγS, where the affinity for ATPγS does not appear to be higher than that for ADP or AMP (**Figure 1-figure supplement 2d**). The lack of preference of RNF213 for ATP provokes the hypothesis that the inhibitory effect of ADP/AMP has a regulatory role. Accordingly, RNF213 might be sensing the relative concentrations of different adenosine nucleotides in the cell, tuning its E3 activity. The energy status of living cells is commonly described by adenylate charge, which ranges in cells from ∼0.7 (lower ATP/AMP ratio) to ∼1.0 (higher ATP/AMP ratio) (*27*). We tested this using ABP-labeling of RNF213 as readout for E3 activity, observing decreasing efficiency of ABP-RNF213 coupling as adenylate charge decreased from 1.0 to 0.7 (**Figure 1f**, **Figure 1-figure supplement 2a**). This points to the possibility that RNF213 activity responds to the changing metabolic state of the cell, a feature not previously described for E3 enzymes.

### Identifying the E3 active site of RNF213

Systematic analysis of various subdomain deletions in RNF213 (**Figure 2a,b**) showed that the N-terminus is dispensable for activity, with both ΔN338 and ΔN1487 mutants remaining active (**Figure 2b**). The C-terminal deletion mutant (ΔCTD, deleted residues 4930-5148) shows a greatly diminished activity, while deletion of ZF2 (ΔZF2, deleted residues 4433-4499), a flexible appendix of the E3 module, is fully inactive. Accordingly, CTD and ZF2 domains are required for E3 activity (**Figure 2b**). These two domains are essential with either UBE2D3 or UBE2L3 (**Figure 2-figure supplement 1**), suggesting that the two E2 conjugating enzymes work with RNF213 in the same way, consistent with previously reported behavior of wild type and ΔRING variants (*16*).

The CTD is a well-structured domain, centrally positioned in the 3D structure of RNF213 (**Figure 2a**), but it does not resemble any known domains and its function is also unknown (*16*). The ZF2 domain is predicted to be a functional zinc finger, highly similar to an uncharacterized zinc finger in the C-terminal region of the human helicase ZNFX1 (**Figure 3c**). ZNFX1 was shown to associate with stress granules (*28*), and is involved in the immune response against both viral (*29*) and bacterial (*28*) infections. Ultraviolet-visible spectroscopy (UV-VIS) revealed that RNF213 ZF2 is indeed a canonical zinc finger (*30, 31*) that can bind divalent transition metals, with a preference for Zn^2+^ (**Figure 3-figure supplement 1b**). Residues Cys4451, His4455, Cys4471, and Cys4474 coordinate the metal ion (**Figure 3a**), as evidenced from altered complex formation of Ala and Ser point mutations (**Figure 3-figure supplement 2b**) (*30, 31*). Metal binding in the ZF2 domain is essential for the E3 ligase activity of RNF213, as mutating these residues renders the protein fully inactive (**Figure 3d**,e). This is not the case for the partially conserved Cys4457, which is involved neither in metal binding, nor in E3 ligase activity (**Figure 3d**, **Figure 3-figure supplement 2b**).

**Figure 3.**
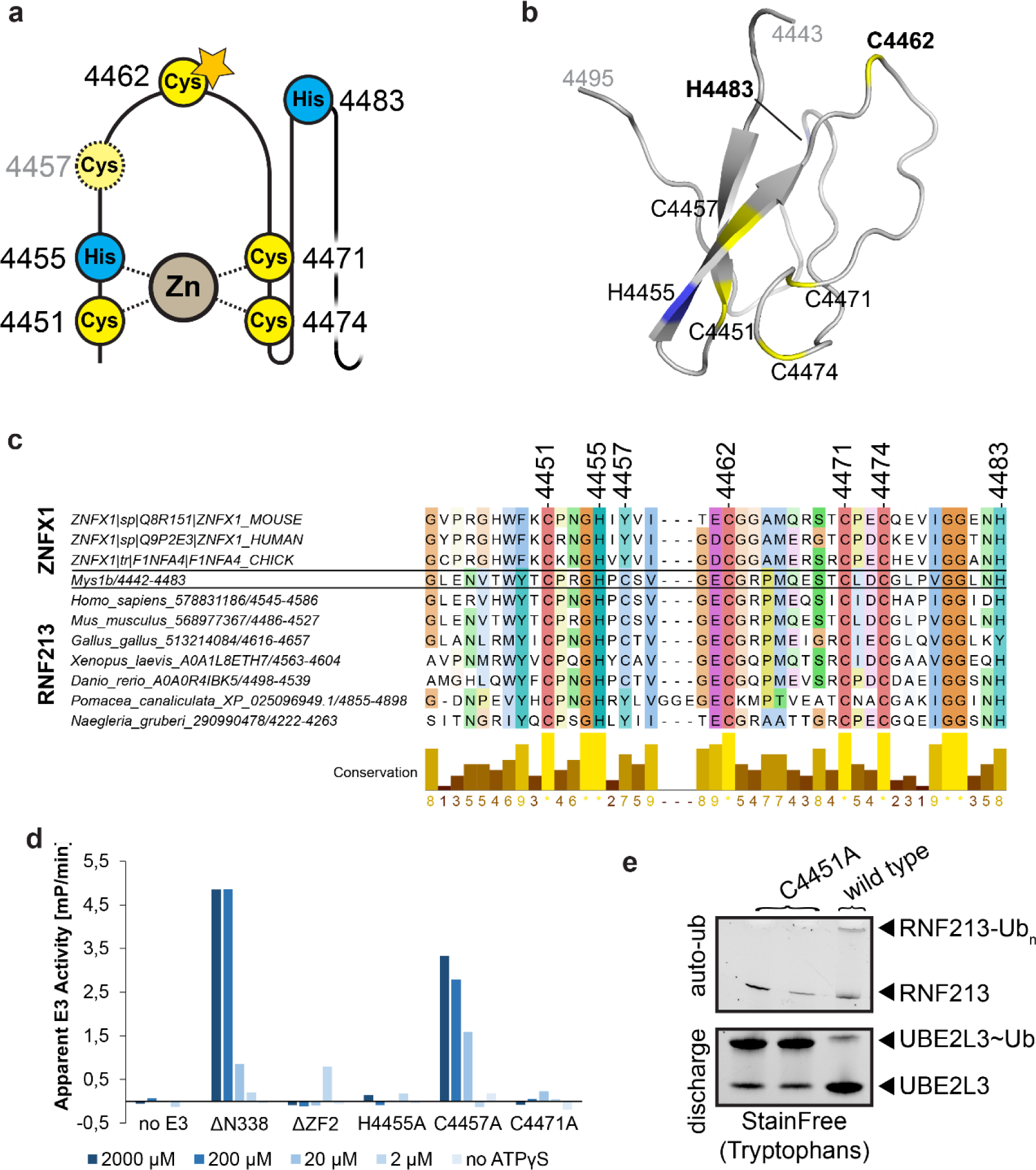
Characterization of the ZF2 domain. (a) A schematic depiction of the zinc finger motif of the ZF2 domain, with the conserved Cys and His residues indicated. Cys4451, His4455, Cys4471, and Cys4474 coordinate the Zn^2+^ ion. Cys4462 is the active site cysteine that accepts the Ub moiety from the E2 enzyme. Poorly conserved Cys4457 is not participating in zinc coordination or fulfilling an enzymatic role. Details about metal binding are provided in Figure 3-figure supplement 1,2. (b) Computational model of the ZF2 zinc finger, with the residues highlighted as in the schematic on the left. More details are provided in Figure 3**- figure supplement 3.** (c) Multiple sequence alignment of representative members of the RNF213 ZF2 domain, and a homologous domain from the antiviral helicase ZNFX1. The residues determined to be involved in Zn binding are fully conserved, as is the active site Cys4462. Jalview conservation score is displayed below the alignment. (d) UBE2L3∼Ub discharge activity of point mutants of the forementioned cysteine and histidine residues, as quantified from fluorescence polarization timecourse curves. Change in polarization over time is used as the measure of apparent activity. Point mutations of the residues involved in zinc binding lead to a complete loss of activity, equivalent to deletion of the entire ZF2 domain. Mutating the poorly conserved Cys4457 has only a minor effect on activity. Four reactions are performed for each construct, with ATPγS concentration between 0 and 2 mM as indicated. (e) SDS-PAGE based UBE2L3 discharge assay with the C4451A mutant. No E2 discharge or auto- ubiquitination can be observed for the mutant, as opposed to the wild type protein which shows strong signal for both. Two batches of the purified C4451A variant are used in lanes 1 and 2.

Apart from the consensus CHCC motif engaged in Zn^2+^ binding, ZF2 contains another strictly conserved cysteine (Cys4462 in mouse, Cys4516 in human RNF213) located on an extended loop (**Figure 3a**,b, **Figure 3-figures supplement 3**). In contrast to all other fully conserved ZF2 residues, mutating Cys4462 (C4462A) does not affect metal binding properties of the zinc finger (**Figure 3-figure supplement 2b**). We thus hypothesized that Cys4462 plays a different, yet mechanistically important role. To test whether Cys4462 is the catalytic cysteine nucleophile involved in E2-E3 transthiolation, we carried out a cross-linking mass spectrometry (XL-MS) analysis of the generated RNF213-ABP adduct (*20, 26*) (**Figure 4a**). In support of our hypothesis, we could unambiguously map the E2:RNF213 cross-links to the ZF2 domain. Importantly, spectral matches converged on Cys4462 (41 spectral matches), with fewer matches to Cys4457 (3 spectral matches) and Cys4451, Cys4471, or Ser4458 (1 spectral match each) (**Figure 4b**). These alternative sites could be ascribed to background labelling that is exacerbated by the high sensitivity and non-quantitative nature of the MS analysis. In sum, our data demonstrate that ZF2 is the primary catalytic element of the RNF213 E3 module, with Cys4462 being the reactive nucleophile participating in the E2-E3 transthiolation reaction.

**Figure 4.**
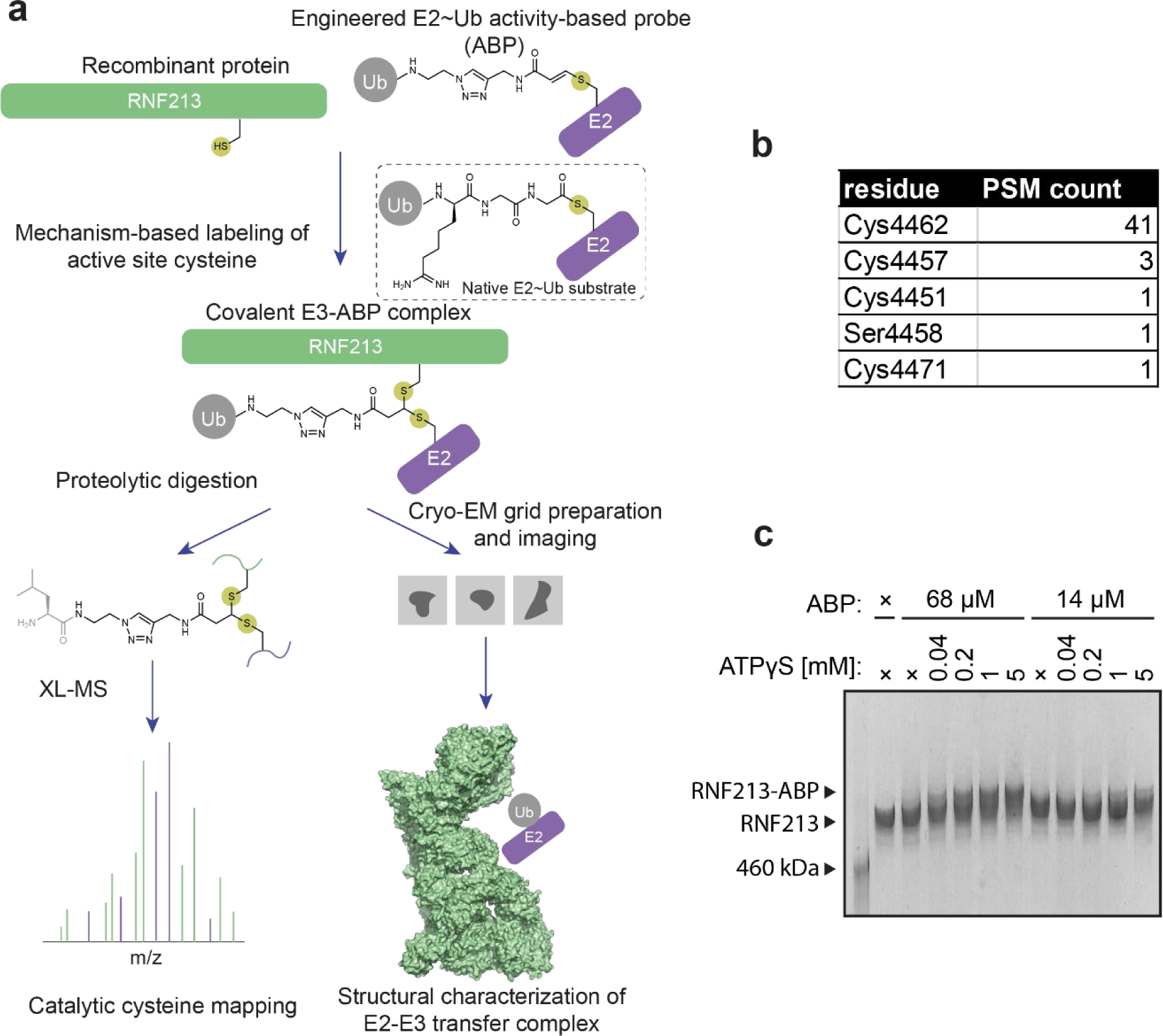
Cross-linking mass spectrometry (XL-MS) analysis of the RNF213-ABP complex. (a) The activity-based probe (ABP) mimics the thioester-linked E2∼Ub intermediate and can be used to produce a stable ternary complex is formed that is representative of the otherwise transient transfer complex. The cysteine in the E3 that is labelled can be identified by proteolytic digestion of the ABP-labelled complex followed by cross-linking mass spectrometry (XL-MS). Structural analysis of the stabilized ternary complex can also be performed, thereby revealing how E2∼Ub is recruited and how transthiolation is mediated. (b) Detected cross-link sites in RNF213-ABP that have an e-value lower than 1·10^-4^ using a false discovery rate (FDR) of 1%. No significant hits are found outside of ZF2. Example MS^2^ spectrum is provided in Figure 4-figure supplement 1. (c) Nucleotide-dependent formation of the RNF213-ABP adduct, analyzed by SDS-PAGE on 3-8% Tris-Acetate gels. A clear shift of the RNF213 band can be observed when the cross-linked adduct to ABP is formed. The gel image has been stretched for a clearer visual appearance.

In order to characterize the active site further, we performed a cryo-EM analysis of mouse RNF213 coupled to the UBE2L3 ABP (**Figure 4a**), a complex mimicking the otherwise transient E2-E3 transthiolation intermediate, and determined its structure at a resolution of 3.5 A (**Figure 5a,b**, **Figure 5-figure supplement 1**). E2 is bound via a well-defined, specific interface with the CTD domain of RNF213. As seen in complexes with other RING, HECT, RBR, and RCR E3 ligases (*32–35*), the interaction site of the E2 is composed of its N-terminal helix and adjacent structural motifs (**Figure 5c**, **Figure 5-figure supplement 2**). Most notably, RNF213 CTD residues Phe5092, Trp5097, and Tyr5106 are properly oriented to undergo hydrophobic interactions with Phe63 located on the L1 loop of UBE2L3 (**Figure 5d**). This contact is strengthened by nearby electrostatic interactions formed between Glu5108(CTD)- Arg6(E2-H1) and Asp5101(CTD)-Lys96(E2-L2). The secondary binding site of the E2, which is formed by a protruding portion (residues 4605-4618) of the RNF213 E3-shell domain is less well defined, comprising mainly non-specific Van der Waals contacts, but possibly helps to properly position the E2 enzyme (**Figure 5e**). Overall, this structural model is in agreement with the demonstrated loss of activity in the ΔCTD mutant (**Figure 2b**). Finally, active site Cys86 of the E2 is oriented towards the center of the E3 pocket, facing a low-contour-level density that is sandwiched between Cys86 and the body of the E3-core domain and presumably originates from ZF2 (**Figure 5-figure supplement 3**). Although the ZF2 density is not well- resolved in the E3:E2 complex (this study) and the apo structure (PDB: 6TAX, 6TAY), the flexible ZF2 domain is located immediately opposite to the E2 Cys86 residue, thereby being properly positioned to participate in the transthiolation reaction. The gap formed between the ABP and well-defined parts of RNF213 is sufficient to accommodate a computed model of the core ZF2 domain, generated with Robetta (*36*) (**Figure 5-figure supplement 3a**). MS analysis confirms that the RNF213-ABP sample contains the Ub moiety, indicating that Ub is still present on the probe as expected, but does not have a well-defined position in the transition state captured by this complex. As previous structural data on UBE2L3-Ub complexes with E3 ligases show that the relative positions of Ub and E2 can vary significantly, the Ub component might be intrinsically mobile (**Figure 5-figure supplement 3c**). Taken together, the E3:E2 complex structure supports our biochemical findings that Cys4462 is the acceptor cysteine of RNF213, equivalent to the active site cysteines present in HECT/RBR/RCR E3 ligases. Thus, RNF213 represents a distinct type of a transthiolation ubiquitin ligase.

**Figure 5.**
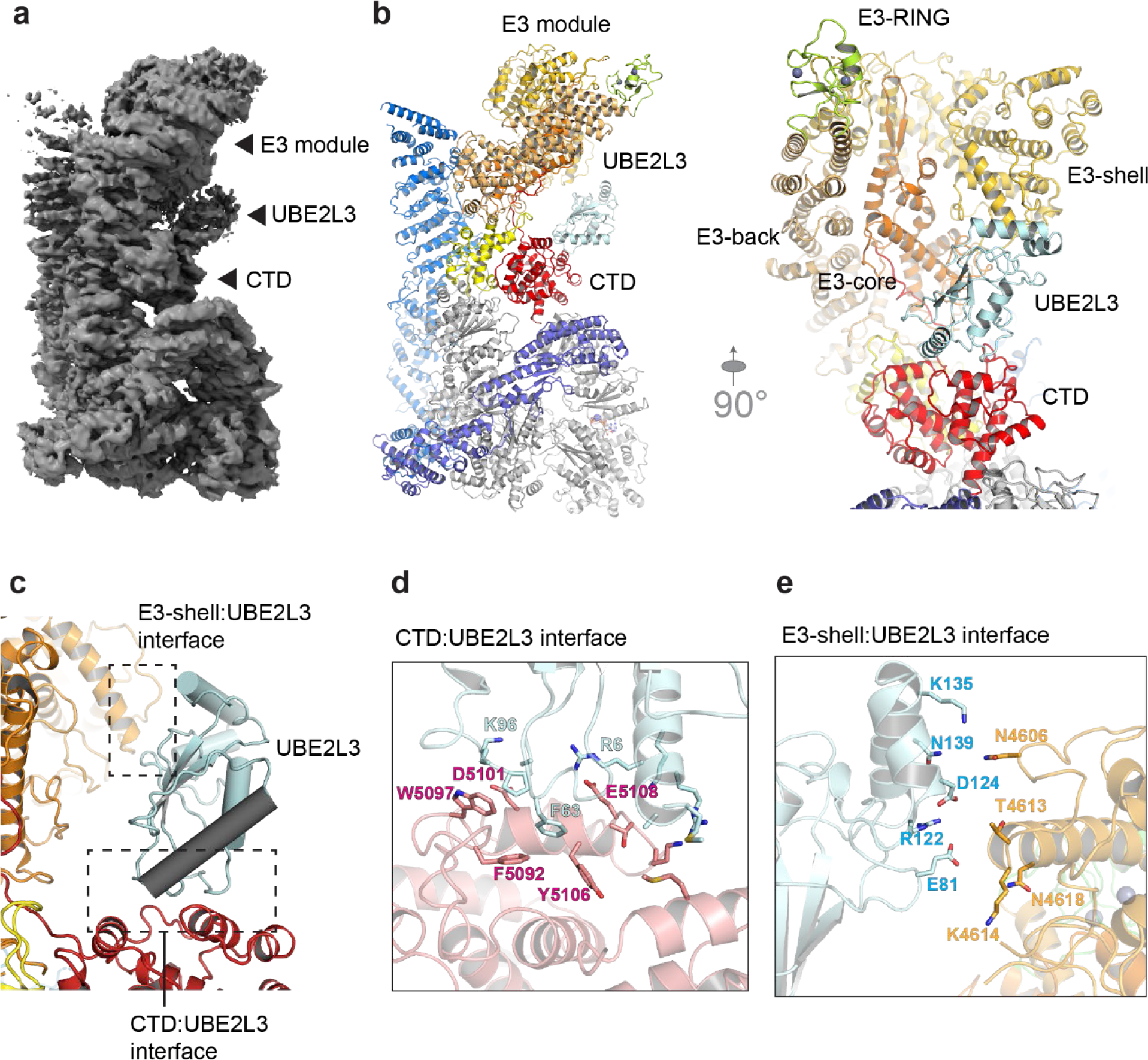
Molecular model of the chemically stabilized RNF213:UBE2L3∼Ub transthiolation intermediate. (a) RNF213-ABP adduct cryo-EM density at 3.5 Å resolution. A distinctive density appears between the E3 shell and the CTD domains of RNF213. Details about the cryo- EM reconstruction are given in Figure 5**-figures supplement 1**. (b) A ribbon model of RNF213-ABP, using the same color code as in Figure 2. Left – side view. Right – front view. UBE2L3 moiety of the UBE2L3∼Ub-anologous ABP can be perfectly fit in the experimental density. The Ub moiety on the ABP and ZF2 of RNF213 are described in Figure 5-figure supplement 3. (c) Zoomed-in view of bound UBE2L3 in the surrounding context of RNF213. Two putative interfaces, with CTD and E3 shell are indicated with dashed boxes. Comparison with E2 interfaces with other E3 enzymes is given in Figure 5-figure supplement 2. (d) Primary E2-RNF213 interface at the CTD domain. (e) Secondary interface with the E3 shell. The distances appear too large to form direct interactions. The side of UBE2L3 near the putative secondary interface is represented by a lower-resolution density, indicating that UBE2L3 is not fully fixed in place.

### Structure of a nucleotide-bound RNF213

With RNF213 being dependent on ATP binding to be an active E3 ligase, the previously reported structures should represent the inactive apo state (PDB: 6TAX, 6TAY). To better understand the switch in activity on a structural level, we aimed to obtain the cryo-EM structure of a nucleotide-bound state of RNF213. For this purpose, we analyzed mouse RNF213 incubated with ATPγS by cryo-EM, reconstructing the EM map at 4.0 Å resolution (**Figure 6a**, **Figure 6-figure supplement 1**) and using it to determine the structure of the nucleotide-bound state. We previously postulated that large conformational changes might occur, changing the relative orientation and positioning of the sub-modules of RNF213. In the present nucleotide-bound state, no such changes were observed. However, the reconstructed map shows clear EM density for the ZF2 domain, which appears to adopt a more defined position upon nucleotide binding (**Figure 6c**). It was possible to fit the computed ZF2 model into the presumed ZF2 density (**Figure 6b**). Moreover, in the ATPase core, nucleotide densities are clearly present in the AAA2, AAA3, and AAA4 active sites, but the exact chemical identity of the nucleotide cannot be unambiguously assigned due to the limited resolution of the map (**Figure 6-figure supplement 2**). Still, the EM density obtained by focused refinement of the AAA core, resolving this region at a resolution of about 3.5 Å, suggests that AAA2 has ATP bound as in the previously determined structures (PDB: 6TAX, 6TAY) (*16*), AAA3 has ADP bound, whereas AAA4 seems to be occupied by a mixture of ADP and ATPγS, with the terminal thiophosphate group being coordinated by Arg3127, the arginine finger of AAA5 (**Figure 6-figure supplement 2**). We presume that nucleotide binding triggers a cascade of small rearrangements in the RNF213 structure, leading to long-range allosteric effects in the E3 module, culminating in the RNF213 ZF2 domain getting fixed in place, thus allowing it to serve as the acceptor site.

**Figure 6.**
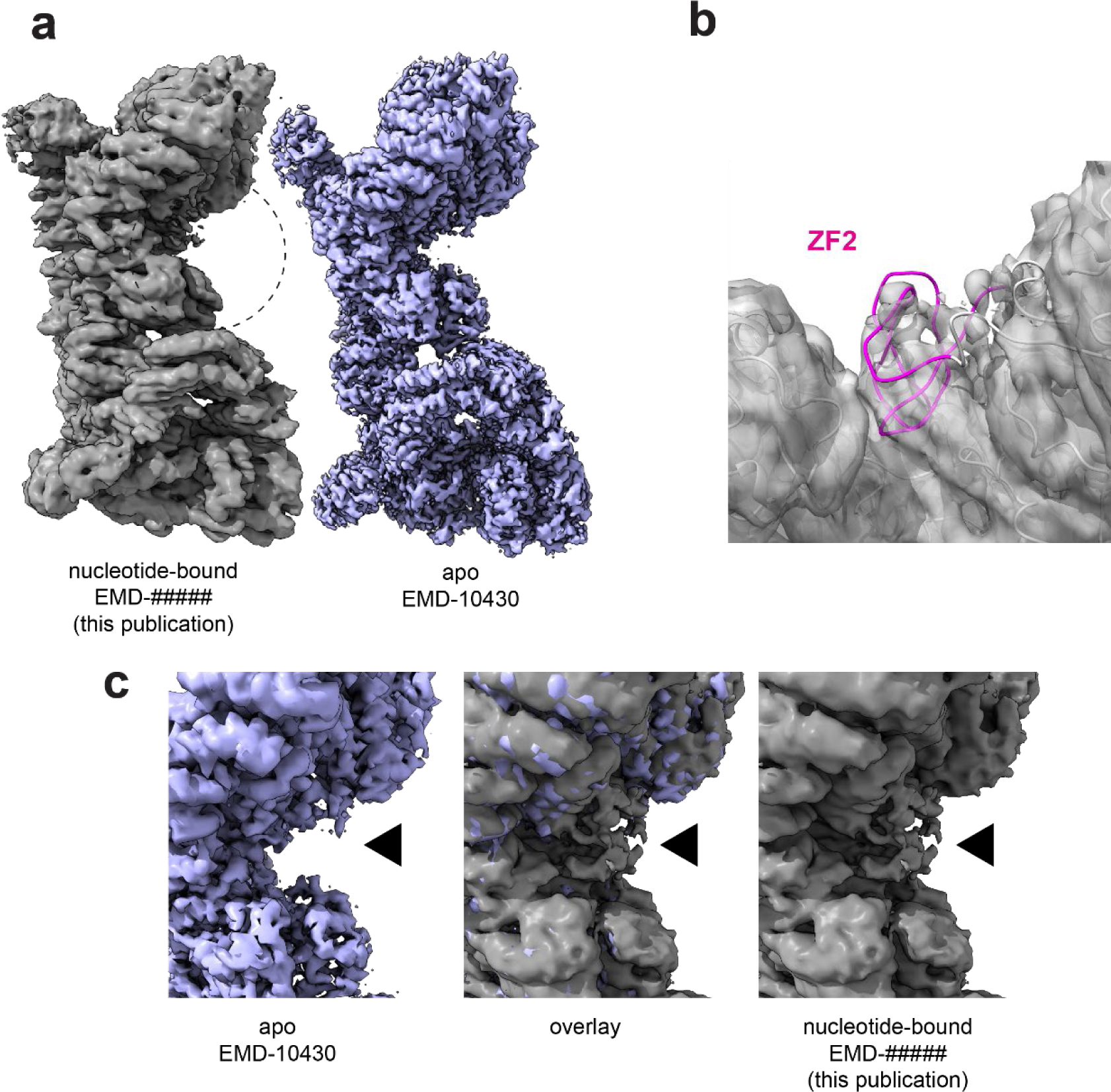
Structure of nucleotide-bound RNF213. (a) Cryo-EM density of nucleotide-bound RNF213 (left), showing no large-scale rearrangements relative to the apo structure (EMDB: EMD-10430, right). The region of interest indicated with a dashed black circle. Details of the cryo-EM reconstruction are given in Figure 6**-figures supplement 1**. (b) Fit of the ZF2 domain into the density appearing in the nucleotide-loaded RNF213. (c) Zoomed-in image of the region between the CTD and E3 shell domains of RNF213, showing the appearance of the density for ZF2. ATPase active sites are analyzed in detail in Figure 6-figure supplement 2.

## Discussion

In the present study we identify and characterize the E3 active site of RNF213, revealing a new type of transthiolation E3 enzyme. We show that RNF213 utilizes its ZF2 domain as the catalytic component for its E3 ligase activity. Once the ubiquitin-loaded UBE2L3 has docked to the CTD, ubiquitin is transferred to Cys4462, the primary acceptor site of RNF213. This first step of the ubiquitination reaction seemingly proceeds in a HECT/RBR-like fashion. Downstream mechanistic steps remain to be resolved, including the question of whether the thioester formed on Cys4462 in ZF2 is used to directly transfer ubiquitin to the substrate. RBR and RCR E3s contain zinc-binding transthiolation domains that can mediate modification of amino and hydroxy substrates (*20*). For these E3 subtypes, the cysteine responsible for substrate modification is in a structured region of their respective zinc-binding domains and in immediate proximity of a free histidine, as seen in MYCBP2 (*20*). The histidine acts as a general base that deprotonates the substrate nucleophile and its mutation typically abolishes activity (*20, 37*). A similar function might be ascribed to the highly conserved His4483 in the RNF213 ZF2 domain (**Figure 3c**). Remarkably, ZF2 models computed with Robetta suggest that His4483 is located on a flexible loop region adjacent to the loop carrying the active site cysteine Cys4462 (**Figure 3-figure supplement 3**). Building on the presented data, follow-up studies can now address ubiquitin’s subsequent route to the substrate.

In contrast to MYCBP2, RNF213 is able to perform lysine ubiquitination, including the formation of polyubiquitin chains (*16*), exemplifying the differences between these two proteins. The identification of ZF2 and Cys4462 as the core E3 elements of RNF213 also raises the possibility of searching for related E3 ligases that share the same motif, similarly to how the RING-type E3s have been annotated. Indeed, residues that are required for RNF213 ZF2 zinc-binding and transthiolation are conserved in the zinc-finger domain in ZNFX1 (**Figure 3c**), raising the possibility that it too is an undiscovered E3 subtype. In support of this, ZNFX1 is enriched in transthiolation activity-based proteomic data relative to controls (*20*). Moreover, the fact that RNF213 was annotated as a RING-type E3 ligase suggests the necessity for caution in such approaches, and highlights the value of biochemical and structural studies of E3 ligases regardless of whether they have already been annotated based on sequence similarity.

Aside from providing first mechanistic insight into its E3 mechanism, our work also demonstrates how the distinctive E3 activity of RNF213 is regulated. We show that ATP binding stimulates the E3 activity, with ADP and AMP acting as competitive inhibitors versus ATP. We also demonstrate that this nucleotide regulation happens already at the step of E2-E3 trans-thiolation that precedes substrate ubiquitination, though we cannot exclude that ATP binding or hydrolysis might additionally affect the downstream activities (*e.g.* substrate modification). Nevertheless, our findings hint towards the possibility of RNF213 sensing the metabolic state of the cell, by modulating its activity based on adenylate charge. Such regulation occurs with the kinase activity of the highly conserved metabolic sensor AMP- activated protein kinase (AMPK), which plays a key role in cellular energy homeostasis (*38*). Our data support the idea that RNF213 functions as metabolic gatekeeper, being involved in fatty acid metabolism (*11*), lipid droplet formation (*12*), and non-mitochondrial oxygen consumption (*13*). Alternatively, nucleotide regulation could work as a proper on-off switch if RNF213 would be found in a local region with very low ATP concentration or adenylate charge. RNF213 was reported to associate with lipid droplets in the cell, where local crowding and layering of bound factors could produce such an environment. Finally, it could be a response to a sudden stressor that changes the energy state of the cell. Of note, RNF213 was reported to be involved in the immune response (*39*) and antiviral resistance, being upregulated by IFN-β and IFN-γ (*4*), and contributing to Rift Valley fever resistance in mice (*9*). This leads to a further appealing interpretation that the spike in the host cell adenylate charge observed during the middle phase of the *Chlamydia* infection cycle (*40*) boosts the activity of RNF213 in a timely manner. This hypothesized association to bacterial infection response is further supported by the finding that RNF213 expression is five-fold increased in macrophages during infection by *M. tuberculosis*, and that the protein is detectable in pull-downs of some of this bacterium’s secretory proteins (*41*). This provides a possible identity for the “environmental factor“ that has been proposed to account for the low penetrance of MMD in compromised individuals (*1*).

The data presented here do not offer an explanation for the role of the conserved RING domain of RNF213. Despite it being a hotspot for MMD-associated mutations, the RING domain is fully dispensable for auto-ubiquitination activity of RNF213. The reported activity of the isolated RNF213 RING domain with UBE2D2 and UBE2N (*22*) contrasts this model and our findings about the non-canonical UBE2L3-mediated activity of RNF213 and rather suggests that RNF213 possesses multiple distinct roles in the cell mediated by either RING or ZF2 domains. It is also conceivable that the described CTD-ZF2 activity acts cooperatively with the RING E3 activity towards a single substrate, as observed with Cullin-RBR E3 complexes (*42*). However, the substrate of RNF213 likewise remains unknown, with the activity of RNF213 here assessed by using auto-ubiquitination as a substrate proxy. For the factors that were found to be necessary for activity, *i.e.* ATP binding, the presence of ZF2 and CTD domains and Cys4462 as catalytic nucleophile, the absence of a distinct substrate in our analyses should not challenge the conclusions made. However, it remains possible that further cellular factors are necessary to induce RNF213 activity in the full biological context, involving the evolutionarily conserved RING domain within the E3 module and the N-terminal stalk that carries an elusive carbohydrate binding domain and the zinc finger ZF1 at the very N-terminus (**Figure 1a**). Alternatively, these domains could be involved in regulation or localization of RNF213, providing a platform for the cell to further modulate its E3 ligase activity.

In sum, we have here obtained a molecular understanding of a novel type of an E3 ligase activity, composed of domains hitherto not implicated with E3 ligases. The present work provides the toolset for further characterizations and manipulations of RNF213, allowing future studies to dissect the biological role of RNF213 and clarify its pathological effect in MMD and related vascular disorders. In addition to categorizing a new type of E3 enzyme, our work suggests an unexplored regulatory principle in ubiquitin biology, the direct coupling of the activity of a ubiquitination enzyme with the metabolic state of the cell.

### Data availability

Atomic coordinates and cryo-EM density maps have been deposited in the Protein Data Bank (PDBe) under accession codes EMD-### resp. ### for the structure of the RNF213-ABP adduct, and EMD-## resp. ## for the stucture of RNF213 with ATPγS. The raw micrographs were submitted to the EMPIAR database (EMPIAR-###, EMPIAR-###). The mass spectrometry proteomics data have been deposited to the ProteomeXchange Consortium via the PRIDE (*43*) partner repository with the dataset identifier PXD#####. All other source data are included in the paper.

## Acknowledgements

We thank all members of the Clausen group for remarks on the manuscript and discussions. The cryo-EM data were collected at the cryo-EM platform of the European Molecular Biology Laboratory in Heidelberg, overseen by Felix Weis, and Diamond Light Source in Oxford. We thank Richard Imre from the Protein Chemistry Facility of IMP/IMBA for assistance with acquiring the data and analysis of the ABP cross-linking and Mathias Madalinski for preparing the synthetic peptides. We also thank Jana Neuhold for cloning the RNF213 expression constructs, and and Thomas Heuser for general assistance with cryo-EM work, both from Vienna BioCenter Core Facilities. We thank Grahame Hardie of University of Dundee for valuable discussion. This work was supported by funding from the European Research Council (ERC) under the European Union’s Horizon 2020 research and innovation programme (AdG 694978), an FFG Headquarter Grant (No 852936) and the Austrian Science Fund (FWF, SFB F 79). S.V. and A.F. are funded by UK Medical Research Council (MC_UU_12016/8) and Biotechnology and Biological Sciences Research Council (BB/ P003982/1). We also acknowledge pharmaceutical companies supporting the University of Dundee Division of Signal Transduction Therapy (Boehringer-Ingelheim, GlaxoSmithKline and Merck KGaA). The IMP is supported by Boehringer Ingelheim.

## Author Contributions

J.A., A.F., S.V., and T.C. designed the experiments and coordinated the project. J.A. performed the biochemical, structural, and bioinformatic analyses and wrote the manuscript with inputs from A.F., D.G., S.V., and T.C. A.F. performed all biochemical experiments involving the ABP. D.G. built the molecular models with participation from T.C. E.R. performed the XL-MS analysis. L.D. and A.L. purified the RNF213 variants. Manuscript was revised with inputs from all authors.

## Materials and methods

Protein sequence analysis, cloning of RNF213 expression constructs, expression and purification of RNF213 protein, preparation of fluorescently labeled ubiquitin, the RNF213 SDS-PAGE-based auto-ubiquitination assay, cryo-EM analyses, and model building were performed as in the associated eLife publication (*16*). Discrepancies and new methods are outlined below.

### E2 discharge assay

The E2 discharge assay was performed using a purified UBE2L3∼Ub thioester intermediate. UBE2L3∼Ub was produced by preparing a reaction mixture consisting of 2.5 μΜ human UBA1, 170 μΜ human UBE2L3, 300 μΜ bovine ubiquitin, and 2 mM ATP, and incubating it at 37°C for 20 min. The reaction mix was optionally supplemented with a 1:200 molar ratio of fluorescently labeled human ubiquitin with an N-terminal overhang containing a cysteine residue, labeled either with DyLight 488 Maleimide or DyLight 800 Maleimide (Thermo Fisher Scientific). The reaction was performed in the ubiquitination buffer (25 mM HEPES, 150 mM NaCl, 10 mM MgCl2, 2 mM TCEP, pH 8.0 @ 25°C). Afterwards, the reaction mixture was supplemented with 20 mΜ imidazole and incubated for 1-5 minutes over Ni-NTA agarose beads (Quiagen). Flowthrough was then collected after spinning the suspension down in a cellulose acetate spin column (Thermo Scientific #69702), separating the His-tagged UBA1 from the other components. Finally, the flowthrough was purified by SEC using a Superdex 75 Increase 3.2/300 column (Cytivia), using the ubiquitination buffer as the elution buffer, separating UBE2L3∼Ub from remaining UBA1, free UBE2L3, free Ub, and ATPγS. UBE2L3∼Ub produced in this way had a concentration between 40 and 80 μΜ and contained only a minor fraction (<10%) of free UBE2L3 as a contaminant. Aliquots were flash-frozen and stored at –70°C. The purified adduct was stable on ice for at least 1 week, and for at least 1 year at –70°C, and had remained fully stable for at least 32 freeze-thaw cycles.

### Fluorescence polarization-based ubiquitination assays

The E3 ligase activity of RNF213 variants was measured using a fluorescence polarization (FP) based assay using 25 mM HEPES, 150 mM NaCl, 10 mM MgCl_2_, 2 mM TCEP (VWR #97064- 848), pH 8.0 as the reaction buffer, supplemented with 10 μΜ of BSA monomer (Sigma Aldrich #A1900). The typical concentrations of the components in the standard assay were as follows: 20 μΜ Ub, 0.5 μM E1, 8 μM E2, 0.05 μM E3. The discharge assay components were as follows: 0.025 μM E3, 20 μM E2∼Ub, 1 mM ATPγS. Ubiquitin source (from bovine erythrocytes) was always spiked with a 1:200 molar ratio of Ubiquitin-DyLight488 (from *E. coli*). Human E2 variants UBE2L3 and UBE2D3 were used, purified from *E. coli*. For E1, human UBA1 purified from *E. coli* was used. A 2× master mix was typically prepared with all the components apart from the screened component. 2 µL of the screened component at 2× final concentration was mixed with the master mix, yielding a 4 μL reaction mixture. 3.5 µL of the resultant mixture was transferred to a 1536-well plate (Greiner Bio-One #782900) and briefly centrifugated. The plates were sealed using a UV-transparent plate seal (Greiner, Merck #Z617571) to prevent evaporation and minimize exposure to atmospheric oxygen. Reactions were monitored by measuring the fluorescence polarization over time using the 485-520-520 fluorescence polarization filter (BMG Labtech) in a PHERAstar FS plate-reader instrument (BMG Labtech). Gain was auto-calibrated at each run to achieve ∼25 % signal saturation in a control well, and was always found to be in the vicinity of 1400-1500, depending on the concentration of fluorescently labeled ubiquitin. Reactions were carried out for 6-12 hours at 30°C. For each curve, the reaction rate was derived from the slope of the earliest curve region where the signal started rising monotonously, after the initial temperature equilibration. This was typically the case in the period 10-30 min after inserting the plate into the instrument.

### Metal-binding assay

Synthetic peptides of the ZF2 domain zinc finger motif were solubilized in the RNF213 buffer (200 mM KCl, 25 mM HEPES, 0.25 mM TCEP, pH 7.2). Binding mixtures contained 200 μΜ of the zinc finger. For screening of different metals, 150 μΜ of each metal was used, or otherwise as indicated in the figure. For assaying metal binding of ZF2 mutants, 300 μΜ of CoCl_2_ was used in all mixtures. Spectra were recorded for all mixtures using a small-volume 10 mm quartz cuvette (Hellma ultra Micro, Sigma Aldrich #Z627062) and a Denovix DS-11 FX+ Spectrophotometer, using no baseline correction, and using water as the blanking solution. The curves were baseline-corrected post hoc by setting the absorbance at 850 nm to zero.

### Cryo-EM data collection and analysis

Both the RNF213-ABP and RNF213-ATPγS samples were recorded on a Titan Krios G3 instrument equipped with a Gatan BioQuantum Energy Filter and a Gatan K3 camera. A single standard data collection with no stage tilt was carried out for RNF213-ABP, while RNF213- ATPγS was recorded in two sessions, one with no stage tilt, and one with a stage tilt of 30°. The two datasets for ATPγS were processed together.

Cryo-EM data processing was carried out in relion 3.1, with particle picking carried out by a combination of cryolo and relion 3D template picking using the model generated by cryolo- picked particles. For the RNF213-ATPγS dataset, all particles were used to generate the final model, only using 2D and 3D classification to select the particles from which the highest resolution could be achieved. For the dataset with RNF213-ABP, 3D classification was additionally used to separate the particles representing ABP-coupled RNF213 (∼2/3 of the particles) from free RNF213 (∼1/3 of the particles).

Images of the cryo-EM density were generated by ChimeraX.

### RNF213 model building

To generate the RNF213-ABP model, RNF213 (PDB: 6TAX) and UBE2L3 (PDB: 4Q5E) were docked into the unsharpened map using Chimera (*44*), and subjected to five cycles of real space refinement in Phenix (1.19.1) (*45*) using reference model restraints and morphing. The resulting model was then further refined using cycles of modelling and refinement with Coot (0.9.5) (*46*) and Phenix Real Space Refine.

For the nucleotide-bound RNF213 structure, the map was sharpened using Phenix AutoSharpen and RNF213 (PDB: 6TAX) was docked in the density using Chimera. Five cycles of real space refinement in Phenix using reference model restraints were performed. The resulting model was then docked into the autosharpened 3.5 Å resolution density obtained from focused refinement of the ATPase module, and was refined using Coot and Phenix Real Space Refine. The ATPase module was then cut and pasted into the model obtained from the overall density, and B factors and validation statistics were calculated using the whole map and Phenix Real Space Refine.

### Generating a model of ZF2

The 3D molecular model of the ZF2 domain was generated by trRosetta (*36*), using the full ZF2 sequence. The model was manually adjusted to account for zinc coordination and to fit into the experimental density.

### Preparation of the UBE2L3 ABP

#### Ub-thioester (Ub-SR)

BL21 cells were transformed with pTBX1-UbΔ74-76-T3C encoding Ub(1–73)-intein, recovered for 1 hour in SOB medium, and used to inoculate 100 ml LB containing 100 ug/mL ampicillin, which was incubated overnight at 37 °C while shaking (220 rpm). 10 mL was used to inoculate 1 L cultures the following day, incubating at 37 °C while shaking (220 rpm). At OD_600_ ∼ 0.4, cells were transferred to 25 °C and after 30 min cultures were induced with 0.3 mM IPTG. After 5 h, cells were harvested and frozen. Cells were thawed in lysis buffer (20 mM Na_2_HPO_4_ pH 7.2, 200 mM NaCl, 1 mM EDTA, Complete EDTA-free protease inhibitor cocktail (one tablet per 50 mL of buffer, Roche) and 0.1 mg/mL DNaseI), sonicated on ice and clarified by centrifugation (16,000 rpm). The clarified lysate was filtered (0.22 μm). An XK 26/60 column was packed with chitin resin (NEB), and equilibrated with lysis buffer. The filtered lysate was loaded on to the chitin column using an Äkta FPLC (1 ml/min). The column was washed with 300 mL wash buffer (20 mM Na_2_HPO_4_ pH 7.2, 200 mM NaCl, 1 mM EDTA), then equilibrated with 60 mL cleavage buffer (wash buffer supplemented with 100 mM MESNA) and the column was rested at 4 °C for 66 h. Cleaved Ub-SR (R is CH_2_CH_2_SO_3_H) was eluted with wash buffer, concentrated to 5 mL and purified by semi-preparative RP-HPLC. Fractions containing Ub-SR were verified by LC-MS (Mw 8417 Da), pooled and lyophilized.

#### Ub-azide (Ub-N_3_)

20 mg Ub-SR was reconstituted in DMSO (200 uL) then H_2_O. (1.8 mL). 344 mg 2- azideoethanamine in 50% (v/v) aqueous DMSO/H_2_O was added to Ub-SR and pH adjusted to ∼9.5 with 35 uL triethylamine (TEA) and briefly vortexed. The reaction was incubated at 30°C for 16 hours, checking progress at 1 hour. Protein was purified by RP-HPLC and fractions containing Ub-N_3_ were verified by LC-MS (Mw 8361 Da), pooled and lyophilised.

#### Ub-TDAE

Ub-N_3_ was conjugated with alkyne-functionalised TDAE by copper-catalyzed azide-alkyne cycloaddition. To 10 mM TDAE in DMSO (200 uL) was added, in this order, pre-prepared 20% DMSO (445 uL), phosphate buffer (100 mM Na_2_HPO_4_/150 mM NaCl, pH 7.5) (50 uL) and 419 uM Ub-N_3_ (3.5 mg in 10% (v/v) DMSO/H_2_O) (250 uL). 30 uL pre-mixed 50 mM Cu(II)SO_4_ (5 uL)/50 mM Tris(3-hydroxypropyltriazolylmethyl) amine (THPTA) (25 uL) was then added to the previously prepared TDAE/Ub-N_3_ solution. 100 mM L-Ascorbic acid (25 uL) was then added to the previously mixed components. The reaction was incubated at 23 °C with shaking (850 rpm) for 15 min. The reaction was subject to semi-preparative RP-HPLC and fractions containing Ub-TDAE were verified by LC-MS (Mw 8624 Da).

#### E2-TDAE-Ub

E2 protein was buffer exchanged on a PD-10 column (Cytvia) equilibrated in degassed phosphate buffer (100 mM Na_2_HPO_4_ pH 7.5, 150 mM NaCl), and concentrated to 15 mg/mL. 5 mg Ub-TDAE was reconstituted in DMSO (50 uL) then H_2_O (450 uL), then mixed with 7.5 mg E2 (500 uL) and incubated at 30 °C for 3 h. ABP assembly was monitored by LC-MS and SDS-PAGE. Reactions were purified by size-exclusion chromatography using a HiLoad 16/600 Superdex 75 pg column (GE Life Sciences) coupled to an ÄKTA Purifier FPLC system (1.0 mL min^−1^, 20 mM Tris pH 7.5, 150 mM NaCl).

#### UBE2L3 (C17S/C137S)

BL21 were transformed with pET15 containing N-terminal Hisx6 tag UBE2L3 (C17S/C137S), Hisx6 tag/StrepII tag UBE2L3 (C17S/C137S), recovered for 1 hour in SOB medium, and used to inoculate 100 ml LB containing 100 ug/mL ampicillin, which was incubated overnight at 37 °C while shaking (220 rpm). 10 mL was used to inoculate 1 L cultures the following day and incubated at 37 °C while shaking (220 rpm). At OD_600_ 0.6, cultures were induced with 0.3 mM IPTG and incubated at 37 °C for 5 h. The cells were harvested by centrifugation and stored at −80 °C. Cells were resuspended in ice cold lysis buffer (50 mM Na_2_HPO_4_ pH 7.5, 150 mM NaCl, 20 mM imidazole, 1 mM TCEP, 1 mg/mL lysozyme, 0.1 mg/mL DNase I, Complete EDTA- free protease inhibitor cocktail (one tablet per 50 mL of buffer, Roche)) – 50 mL lysis buffer per litre culture – and incubated on ice for 30 min followed by sonication. The clarified lysate was then subjected to Ni-NTA affinity chromatography (Qiagen) and washed with wash buffer (Na_2_HPO_4_ pH 7.5, 150 mM NaCl, 20 mM imidazole). Protein was eluted with elution buffer (Na_2_HPO_4_ pH 7.5, 150 mM NaCl, 300 mM imidazole), E2-containing fractions confirmed by SDS-PAGE and Coomassie staining, pooled, concentrated, and protein further purified by size-exclusion chromatography using a HiLoad 16/600 Superdex 75 pg column (GE Life Sciences), 1.0 mL/min in Na_2_HPO_4_ pH 7.5, 150 mM NaCl, 1 mM TCEP. Fractions were collected, concentrated and flash frozen for storage at −80 °C. For StrepII-tagged probe, a strepII sequence was inserted between the E2 start codon and the precision protease site. Following binding to Ni-NTA, the Hisx6 tag was cleaved with precision protease (in-house) and StrepII-UBE2L3 (C17S/C137S) eluted.

### Adenylate charge

The adenylate charge (*47*) represents the relative concentrations of ATP and ADP, against the total adenine nucleotide pool (ATP, ADP and AMP):

Adenylate charge = (ATP + ½ADP)/(ATP + ADP + AMP)

In resting cells, the adenylate charge approximates to 1. Upon ATP consumption, ADP and AMP levels increase. Keeping the total adenine nucleotide pool at 1 mM, we incrementally decreased the adenylate charge from 1 to 0.7 (*48*). 1.5 μM RNF213 (ΔN338) was incubated with 68 μM UBE2L3 ABP in the presence or absence of varying concentrations of nucleotides ATPγS, ADP or AMP, in 50 mM Tris.HCl pH 7.5, 2.5 mM MgCl_2_, 150 mM NaCl. Reactions were incubated at 30°C for 4 hours before resolving on 3-8% Tris-Acetate gels.

### Preparation of RNF213-ABP

A 600 μL reaction containing 1.5 μM full-length RNF213 (528 μg), 100 μM StrepII-H7 probe (1.56 mg) and 5 mM ATPγS were incubated in 50 mM Tris.HCl pH 7.5, 2.5 mM MgCl2, 150 mM NaCl at 30 °C for 4 h, with agitation (300 rpm). 1 μL of reaction before and after incubation were assessed by SDS-PAGE on a 3-8% Tris-Acetate gel.

The assembled RNF213-ABP adduct was further purified by SEC using a Superose 6 Increase 3.2/300 column (Cytivia) equilibrated with a buffer containing 200 mM KCl, 40 mM HEPES, 0.5 mM TCEP, pH 8.0. Fractions containing the adduct were identified by SDS-PAGE using a 4-20 % Tris-Glycine gel (BioRad).

### XL-MS analysis of RNF213-ABP

#### Sample preparation

40 ul of the SEC-purified RNF213-ABP sample (0.3 mg/mL, buffer: 200 mM KCl, 25 mM HEPES, 2 mM TCEP, 10 mM MgCl2) was supplemented with urea to a final concentration of 8 M. Dithiothreitol (DTT) was added to 10 mM final concentration and the sample was incubated for 1 hour at 37°C. The alkylation was performed by adding iodoacetamide (IAA) to a final concentration of 20 mM and incubating for 30 minutes protected from light. The reaction was quenched by addition of DTT to 5 mM and incubated again 30 minutes at room temperature.

Then the sample was split in 2 equal parts:

The first part was diluted to 6 M urea with 100 mM ammonium bicarbonate (ABC) followed by addition of 500 ng Lys-C (FUJIFILM Wako Pure Chemical Corporation) and incubation at 37°C for 2 hours. The sample was diluted to 2 M urea with 100 mM ABC, 500 ng trypsin (Promega, Trypsin Gold) were added and incubated at 37°C overnight. The next day the sample was further diluted to 1 M urea and 750 ng Glu-C (Promega, V1651) were added and incubated for another 8 hours at 37°C.

The second part was diluted to 1 M urea with 100mM ABC, then 750 ng Glu-C (Promega, V1651) were added and incubated overnight at 37°C. The next day 750 ng trypsin were added, and the sample was incubated for 8h at 37°C.

Both samples were acidified with trifluoroacetic acid (TFA, Pierce) to a final concentration of 1%. 20% of each sample was analysed by LC-MS/MS.

#### LC-MS/MS on Exploris 480

Generated peptides were analysed on an UltiMate 3000 RSLCnano system, which was coupled via a Nanospray Flex ion source to an Orbitrap Exploris 480 mass spectrometer equipped with a FAIMS pro interface (Thermo Fisher Scientific).

Peptides were loaded onto a trap column (PepMap C18, 5 mm × 300 μm ID, 5 μm particles, 100 Å pore size) at a flow rate of 25 μL/min using 0.1% TFA as mobile phase. After 10 min, the trap column was switched in line with the analytical column (PepMap C18, 500 mm × 75 μm ID, 2 μm, 100 Å, Thermo Fisher Scientific), which was operated at a flowrate of 230 nL/min at 30°C. For separation a solvent gradient was applied, starting with 98% buffer A (water/formic acid, 99.9/0.1, v/v) and 2% buffer B (water/acetonitrile/formic acid, 19.92/80/0.08, v/v/v), followed by an increase to 35% B over the next 60 min, followed by a steep gradient to 90% B in 5 min, staying there for five min and decreasing to 2% B in another 5 min.

The Orbitrap Exploris 480 mass spectrometer was operated in data-dependent mode, performing a full scan (m/z range 375-1500, resolution 60,000, target value 3E6) at 3 different CVs (-45, -55, -65), followed by MS/MS scans to result in a cycle time of 1 second per CV. MS/MS spectra were acquired using a stepped normalized collision energy of 28+/-2%, isolation width of 1.4 m/z, resolution of 30.000, maximum fill time of 100ms, the target value of 2E5 and intensity threshold of 5E3 and fixed first mass of m/z=120. Precursor ions selected for fragmentation (exclude charge state 1, 2, >8) were excluded for 30 s. The peptide match feature was set to preferred and the exclude isotopes feature was enabled.

#### Data Analysis

Raw files were analyzed using pLink 2.3.9 software. All MS/MS spectra were searched against a custom database containing only the 3 proteins Mys1b (based on mouse RNF213), ABP_UBE2L3 (based on human UBE2L3), and Ub-ABP (based on human ubiquitin). The peptide and fragment mass tolerance was set to ±10 ppm. The minimum peptide length was set to 6 amino acids, the maximum peptide mass to 10000 Dalton, the maximal number of missed cleavages was set to 3, and using combined trypsin and GluC as enzyme specificity (both without restriction at proline). ABP was defined as a cross-linker, with a mass of 306.18042 and a composition of H(22)C(14)N(6)O(2) on C,S,T. The same mass was searched as a Monolink mass.

Carbamidomethylation on cysteine, oxidation on methionine, and deamidation on asparagine and glutamine were set as variable modifications. Data were filtered to 1% FDR by pLink and additionally to an E value<0.0001 on CSM level.

---

**Table 1:** Cryo-EM data collection, refinement and validation statistics

|  | RNF213-ABP<br>(EMDB: #####)<br>(PDB: #####) | RNF213 + ATPγS<br>(EMDB: #####)<br>(PDB: #####) |
| --- | --- | --- |
| **Data collection and processing** |  |  |
| Voltage (kV) | 300 | 300 |
| Electron exposure (e<sup>-</sup>/Å<sup>2</sup>) | 32.5 | 45.0 |
| Defocus range (μm) | -0.8 to -2.5 | -1.5 to -2.5 |
| Pixel size (Å) | 1.072 | 1.053 |
| Symmetry imposed | C1 | C1 |
| Initial particle images (no.) |  |  |
| Final particle images (no.) | 98 000 | 336 000 |
| Map resolution (Å) | 3.5 | 4.0 |
| FSC threshold | 0.143 | 0.143 |
| **Refinement** |  |  |
| Initial model used (PDB code) | 6TAX, 4Q5E | 6TAX |
| Model resolution (Å) | 4.2 | 4.3 |
| FSC threshold | 0.5 | 0.5 |
| Map sharpening *B* factor (Å<sup>2</sup>) | 0 | -140 |
| Model composition |  |  |
| Non-hydrogen atoms | 36779 | 35597 |
| Protein residues | 4579 | 4426 |
| Ligands | ATP, Zn (2), Mg | ATP, ADP, AGS,<br>Zn (2), Mg |
| *B* factors (Å<sup>2</sup>) |  |  |
| Protein | 256 | 73 |
| Ligand | 195 | 58 |
| R.m.s. deviations |  |  |
| Bond lengths (Å) | 0.006 | 0.006 |
| Bond angles (°) | 1.32 | 1.32 |
| Validation |  |  |
| MolProbity score | 1.8 | 1.8 |
| Clashscore | 5.4 | 5.2 |
| Poor rotamers (%) | 0.0 | 0.1 |
| Ramachandran plot |  |  |
| Favored (%) | 90.9 | 91.7 |
| Allowed (%) | 8.9 | 8.2 |
| Disallowed (%) | 0.2 | 0.2 |

**Figure 1-figure supplement 1.**
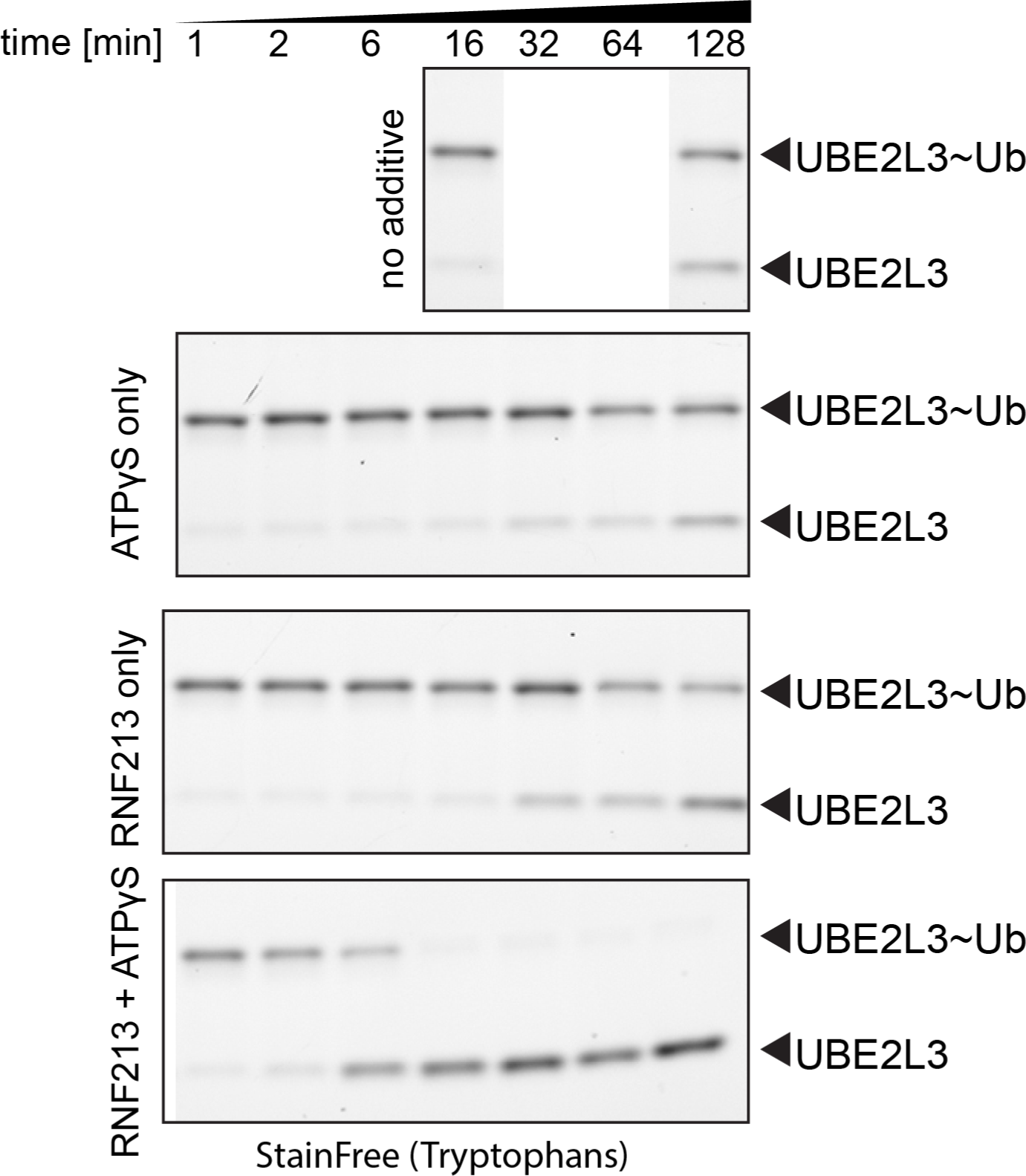
RNF213-mediated discharge of Ub from purified E2∼Ub in presence or absence of ATPγS. Rows from top to bottom are timecourses of UBE2L3∼Ub discharge in absence of either RNF213 or ATPγS, presence of ATPγS only, presence of RNF213 only, and presence of both species, as indicated on the figure. RNF213 shows negligible activity when no nucleotide is present in the reaction, but is strongly activated upon addition of ATPγS.

**Figure 1-figure supplement 2.**
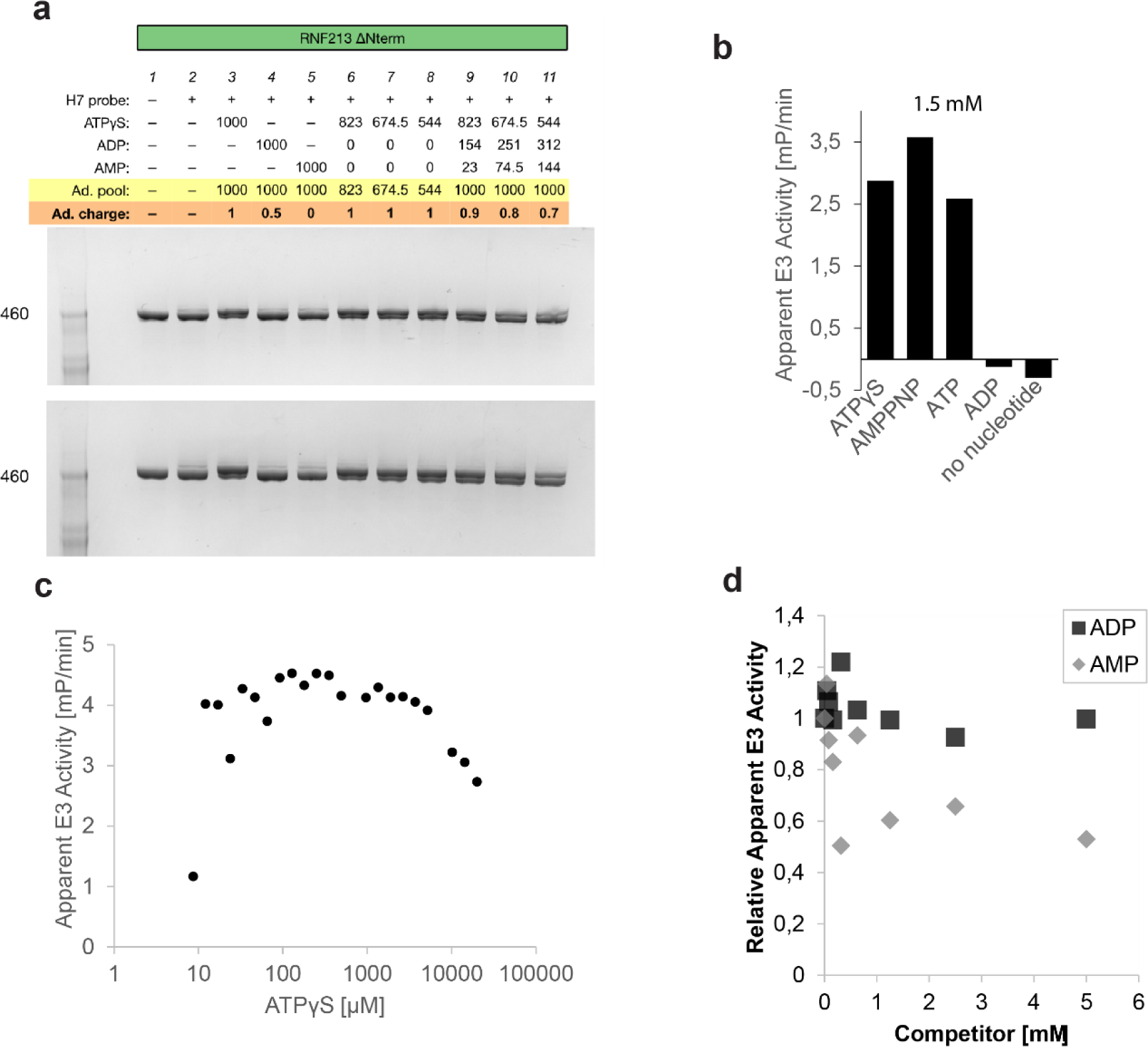
(a) Labeling of RNF213 ΔN338 with the UBE2L3 ABP in conditions mimicking different adenylate charge states in the cell. Each reaction contains 1.5 μΜ RNF213 and 68 μΜ ABP. Two independent experiments are shown. The quantification for these data is given in Figure 1. (b) Quantification of the discharge rate with ATP analogues and ADP. Rates have been determined from Fluorescence Polarization timecourse curves, calculating the slope of the curve after reaching steady state conditions and constant reaction temperature. ATP analogues strongly stimulate RNF213 activity, while ADP stimulates RNF213 only very weakly. (c) Dependence of UBE2L3∼Ub discharge rate on the concentration of ATPγS. Concentrations as low as 50 μM already lead to plateauing of the reaction rate. When ATPγS concentration is very high (>2-5 mM), the reaction rate starts falling off, likely due to non-specific effects. (d) Competition of ADP or AMP with ATPγS in the UBE2L3∼Ub discharge assay. RNF213 is mildly inhibited by ADP, and more strongly inhibited by AMP.

**Figure 1-figure supplement 3.**
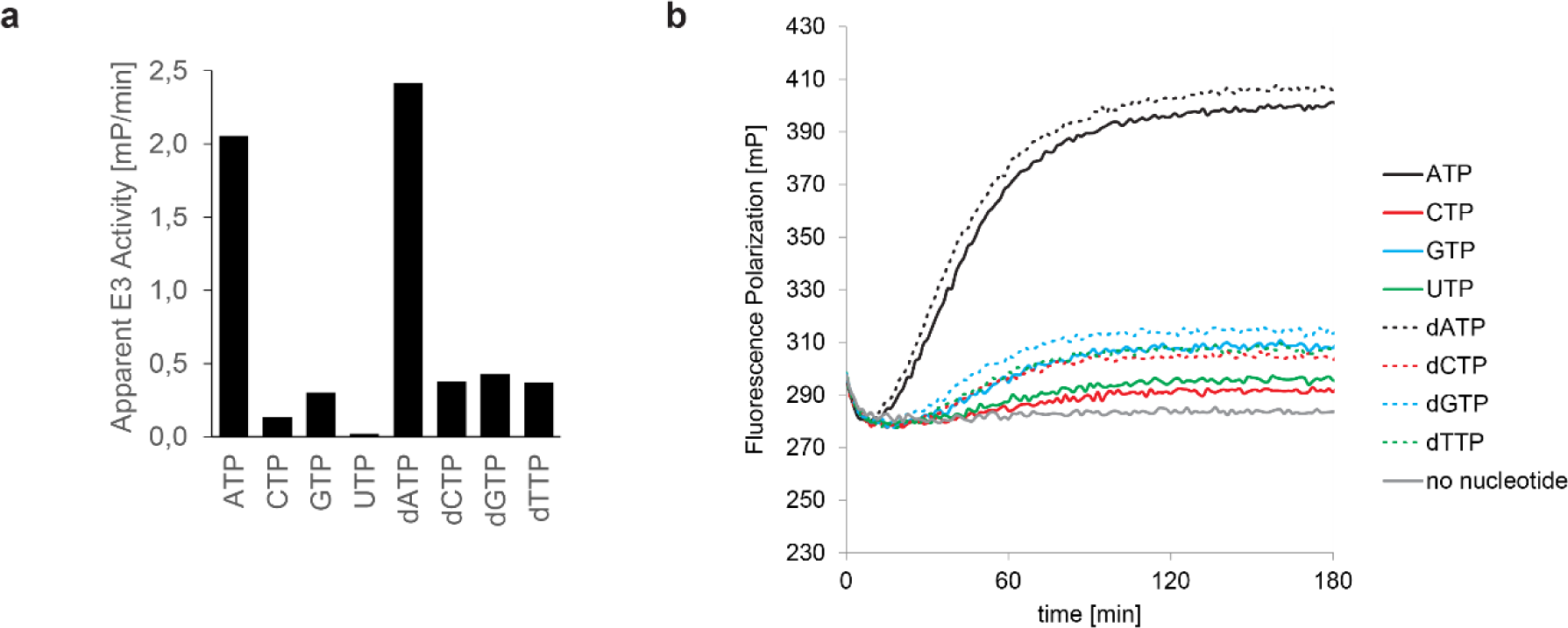
RNF213 activity with non-adenosine nucleotides. (a) Reaction rates with all four canonical NTPs and dNTPs. 20 nΜ RNF213 was mixed with 10 μΜ UBE2L3∼Ub and 0.2 mM of a nucleotide. Only ATP and dATP can efficiently support RNF213 activity. dATP is as efficient as ATP. Change in polarization over time is used as the measure of apparent activity. (b) Fluorescence polarization timecourse curves from which the rate was calculated. A low level of activity is observable with all nucleotides, but only ADP and dADP strongly activate RNF213.

**Figure 1-figure supplement 4.**
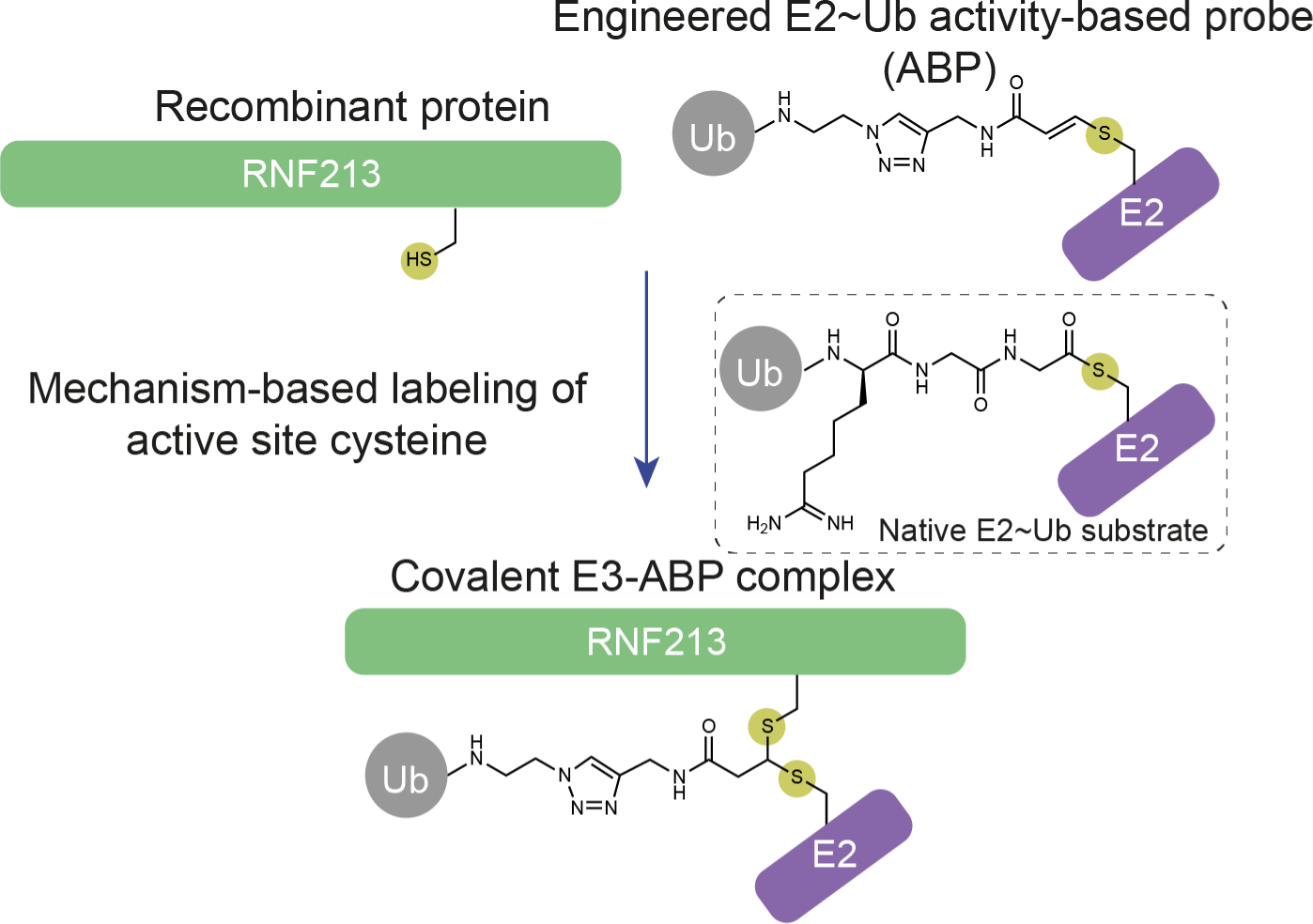
The activity-based probe (ABP) mimics the thioester-linked E2∼Ub intermediate. E3s that utilize an active site cysteine undergo E2-E3 transthiolation, which involves proximity-enabled transfer of the Ub molecule to the E3 active site cysteine. With the ABP, the thioester bond in E2∼Ub has been replaced with a reactive group that traps the E3 active site cysteine. A stable ternary complex is formed that is representative of the otherwise transient transfer complex.

**Figure 2-figure supplement 1.**
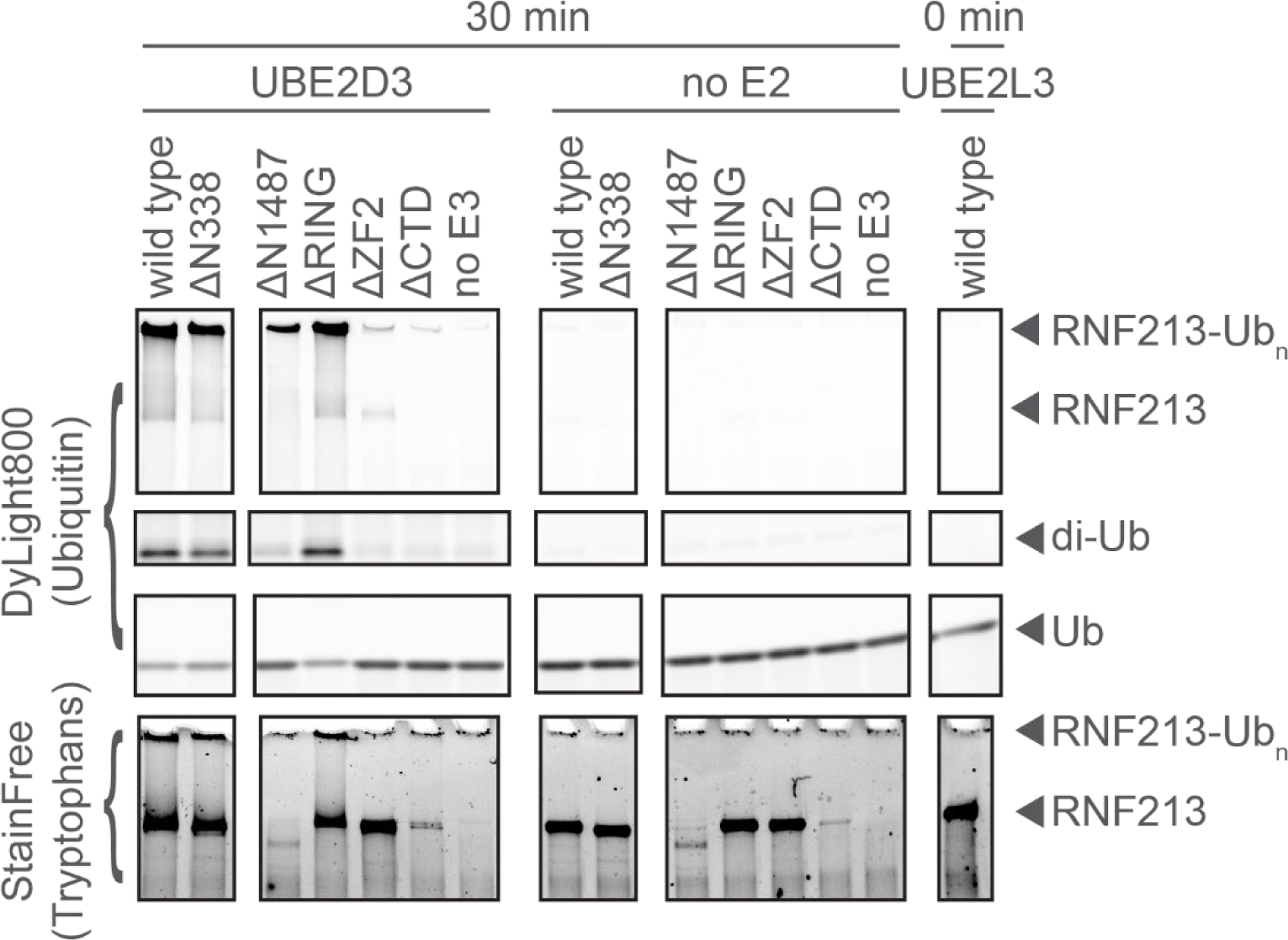
Activity of RNF213 domain deletion mutants with UBE2D3. The promiscuous E2 conjugating enzyme UBE2D3 shows the same activity profile with RNF213 deletion mutants as the preferred E2 UBE2L3 (shown in Figure 2), indicating that UBE2D3 depends on the same mechanism as UBE2L3 to carry out its function.

**Figure 3-figure supplement 1.**
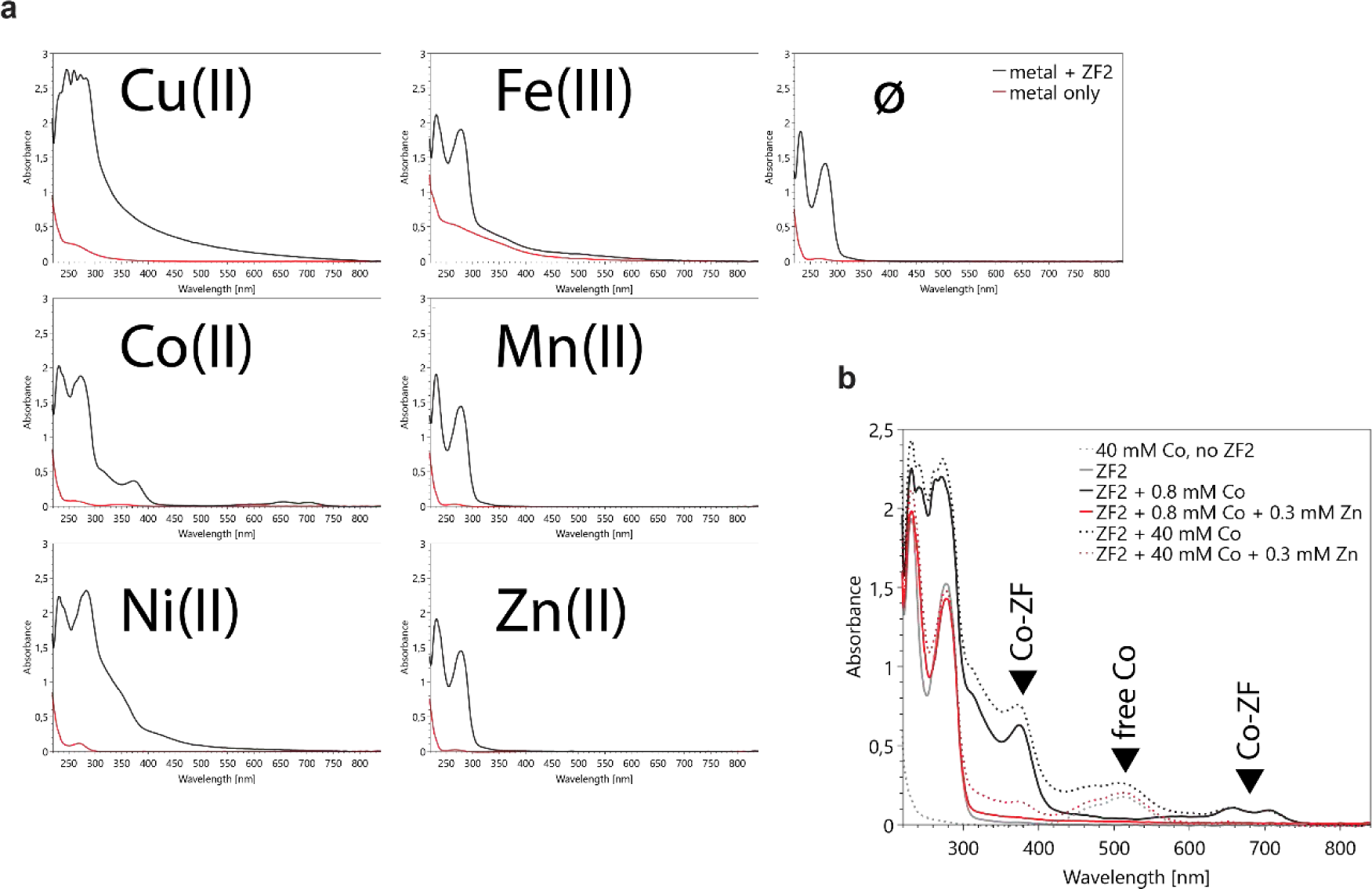
Binding of various metal ions to the ZF2 zinc finger. (a) A synthetic peptide corresponding to the RNF213 ZF2 domain zinc finger is able to bind divalent metal ions of Cu(II), Co(II), and Ni(II), but does not bind the trivalent Fe(III). Binding of Mn(II) and Zn(II) cannot be directly assessed as they do not absorb in the measured UV-VIS wavelength range. (b) Zinc(II) can out-compete bound Co(II) even when a >100-fold excess of Co(II) over Zn(II) is present. The Co(II)-specific spectral signal is greatly diminished, indicating a strong preference of RNF213 ZF2 for zinc over Co(II). Peaks of interest are indicated with a black triangle.

**Figure 3-figure supplement 2.**
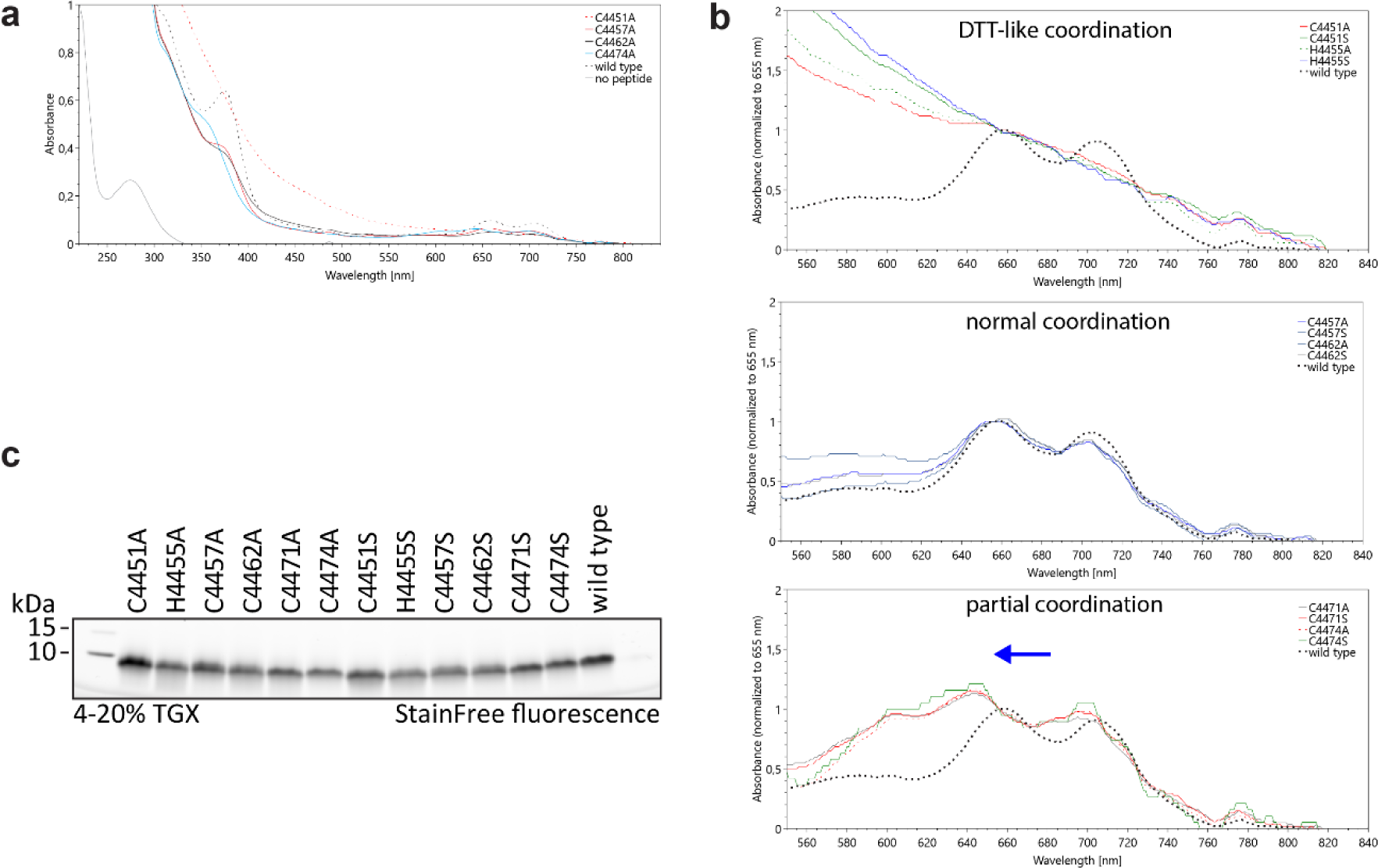
Effect of ZF2 mutations on metal binding. Alanine and serine point mutants of all residues possibly involved in metal binding in the RNF213 ZF2 zinc finger motif were synthetized and their solutions were prepared in the RNF213 buffer at 200 μΜ with a 1.5-fold excess of CoCl2. (a) Full UV-VIS spectra of the formed cobalt complexes of RNF213 ZF2 alanine mutants of all sites. Altered spectra can be observed for C4451A, H4455A, C4471A, and C4474A, while C4457A and C4462A appear similar to the wild type. (b) Detailed view of the spectra of the ZF2-Co complexes, zoomed in on the characteristic Co-zinc-finger complex peak between 600 nm and 800 nm. Three groups of mutations can be observed. Mutations of Cys4451 and His4455 lead to a complete loss of the characteristic peak and to a Co-DTT-like spectrum, giving the solutions a slightly brown tinge. Mutations of Cys4457 and Cys4462 match the spectrum of the wild type. Finally, mutations of Cys4471 or Cys4474 retain the specific Co-zinc-finger peak, but with a clear blueshift, consistent with incomplete coordination. The alanine and serine mutants of each residue match perfectly, making it highly unlikely that the observed effects are non-specific. (c) SDS-PAGE analysis of the peptides used in the metal binding assay.

**Figure 3-figure supplement 3.**
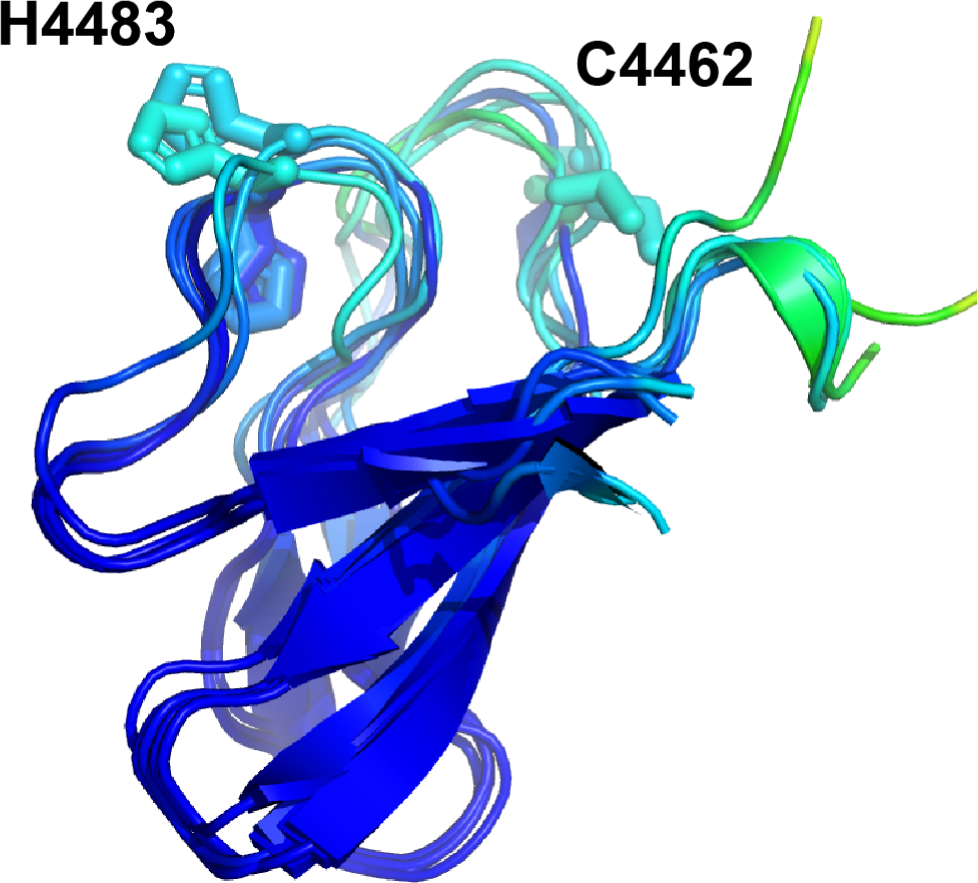
Superposition of different Robetta models of the ZF2 domain, colored by the prediction confidence score. Cys4462 is isolated from the zinc binding site and is exposed at the tip of a loop. A neighboring loop contains His4483, which could work in conjunction with Cys4462, fulfilling the role of the general base that deprotonates amine or hydroxyl Ub acceptors. The low confidence score suggests that both of the loops are flexible.

**Figure 4-figure supplement 1.**
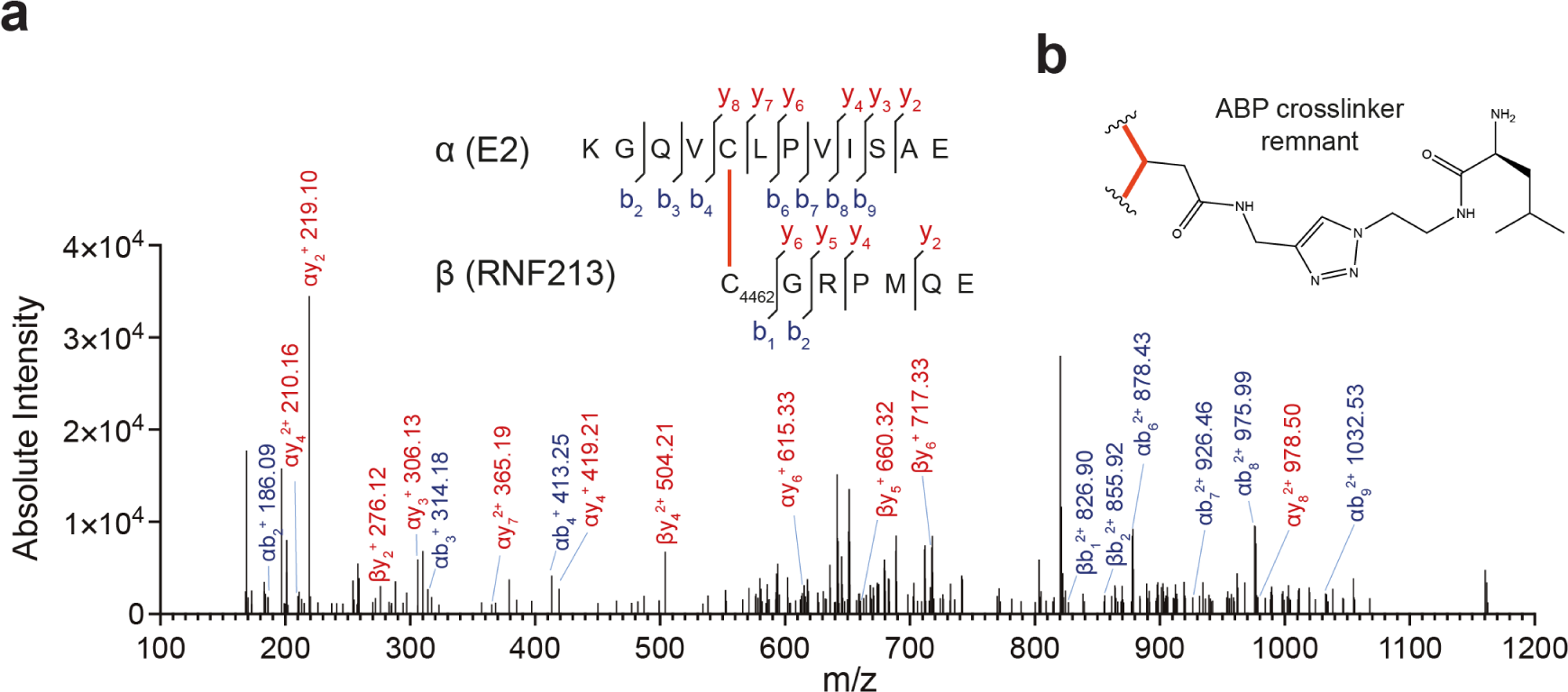
(a) Example MS^2^ spectrum for a cross-linked peptide in which RNF213 Cys4462 is coupled to UBE2L3 Cys86 by the ABP-derived cross-linker. The peptide is obtained from a double digestion of ABP-labeled full-length RNF213 with the endopeptidases GluC and trypsin. The displayed spectrum is for a 4+ precursor ion. Observed *m/z* = 2369.186177; theoretical *m/z* = 2369.188539. (b) Illustration of the cross-linker as it exists in the adduct.

**Figure 5-figure supplement 1.**
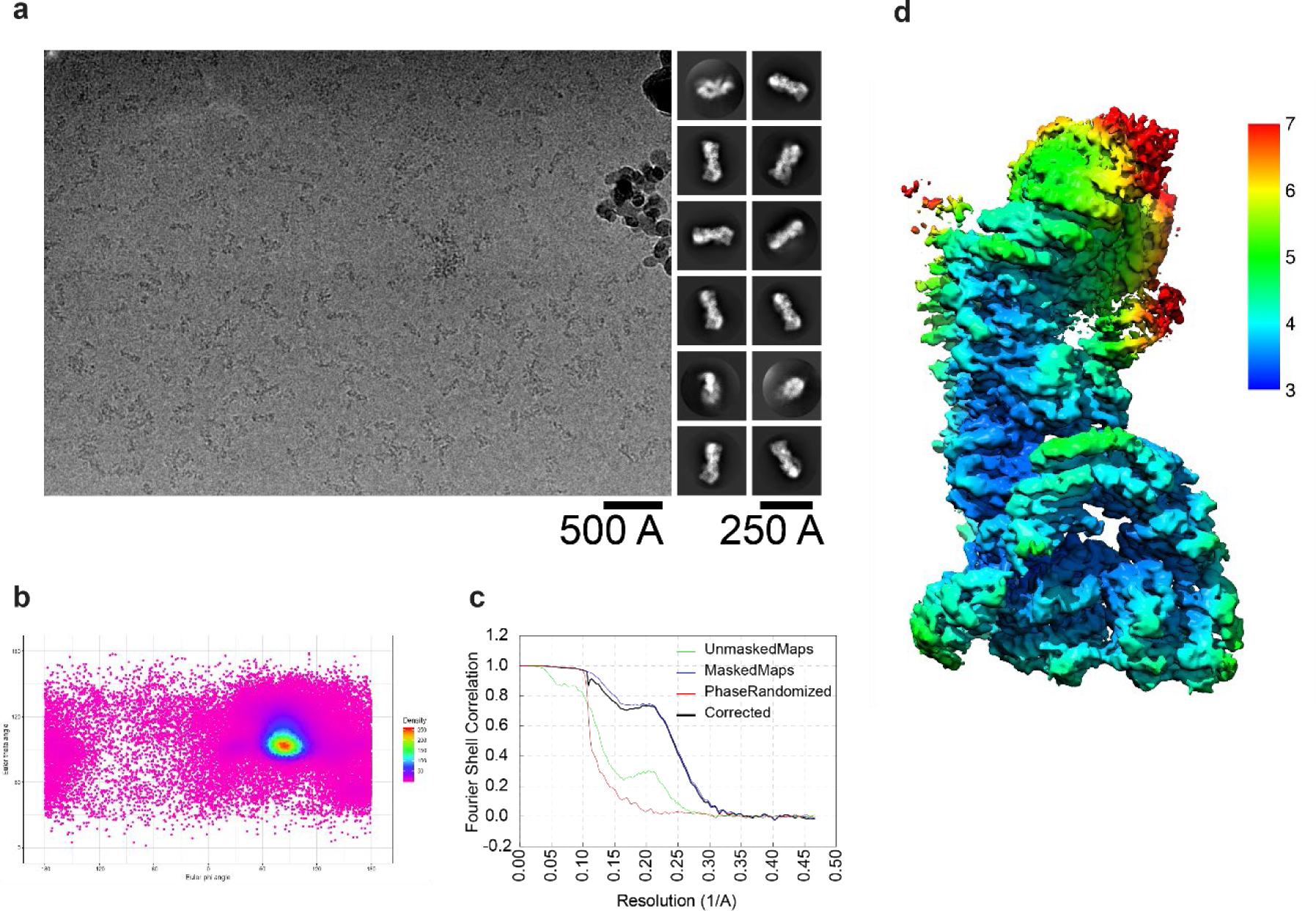
Cryo-EM analysis of the chemically stabilized RNF213:UBE2L3∼Ub transthiolation intermediate. (a) Left: Example micrograph, low-pass filtered at 20 Å for visual clarity. Right: Image class averages as generated by 2D classification in relion. (b) Angular distribution heat map of particles used to reconstruct the cryo-EM density. (c) Fourier Shell Correlation (FSC) curves indicating a resolution of 3.5 Å using the FSC=0.143 criterion. (d) Reconstructed cryo-EM density of the RNF213:UBE2L3∼Ub complex colored by local resolution. The resolution color scale is from 3 Å (blue) to 7 Å (red).

**Figure 5-figure supplement 2.**
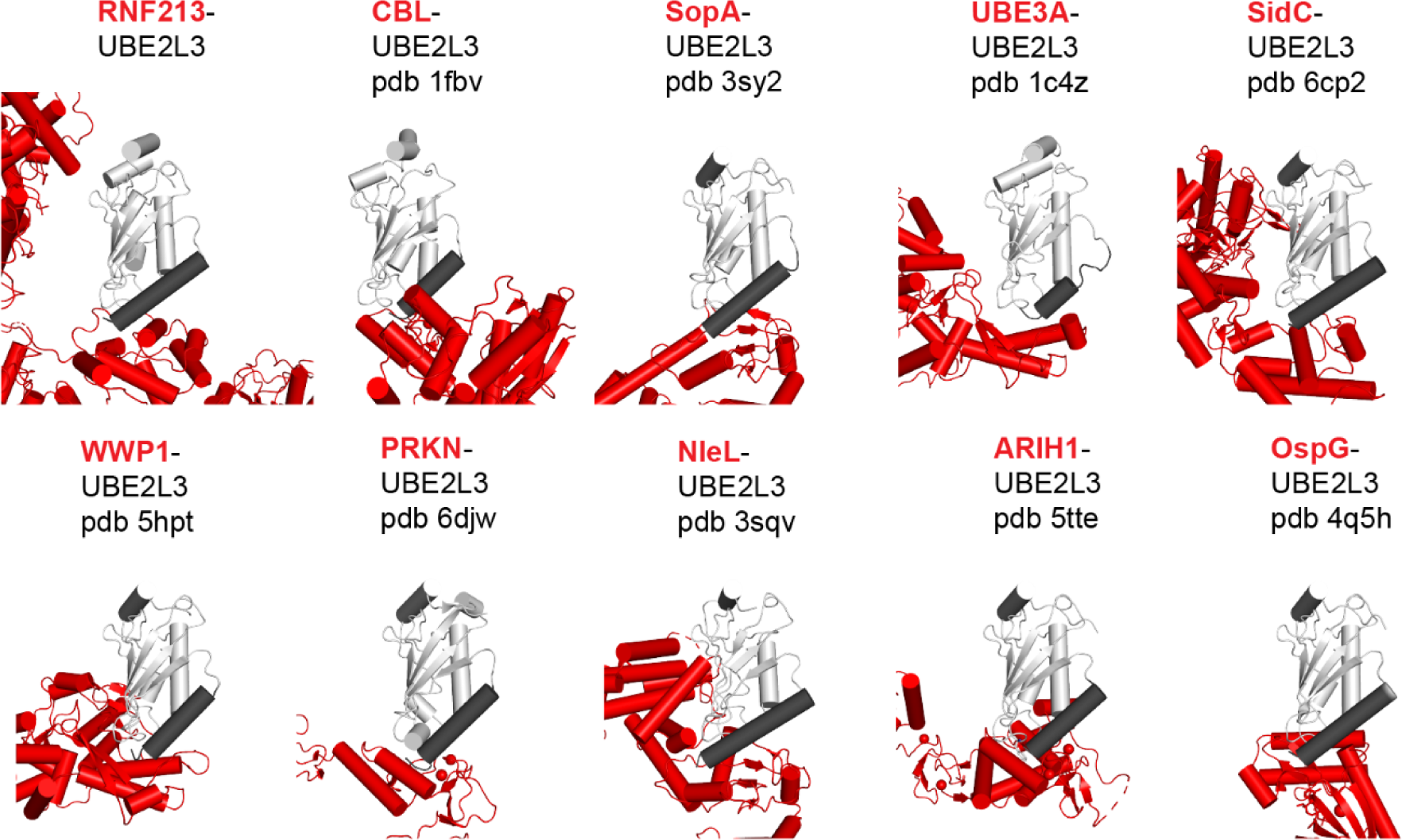
Structural variation in published molecular models of UBE2L3-E3 complexes. Each panel shows a published structure involving an interaction between UBE2L3 (light grey) and an E3 ligase (red). Helix H1 of UBE2L3 in each structure is highlighted (dark grey). Interface between RNF213 CTD and UBE2L3 matches the interface found in all previously known UBE2L3-E3 structures. The secondary E3-UBE2L3 interface is not found in any of the known structures, apart from the non-canonical bacterial E3s NleL and SidC, in which there exists a weak secondary interface that is similar to what appears to be the case for RNF213.

**Figure 5-figure supplement 3.**
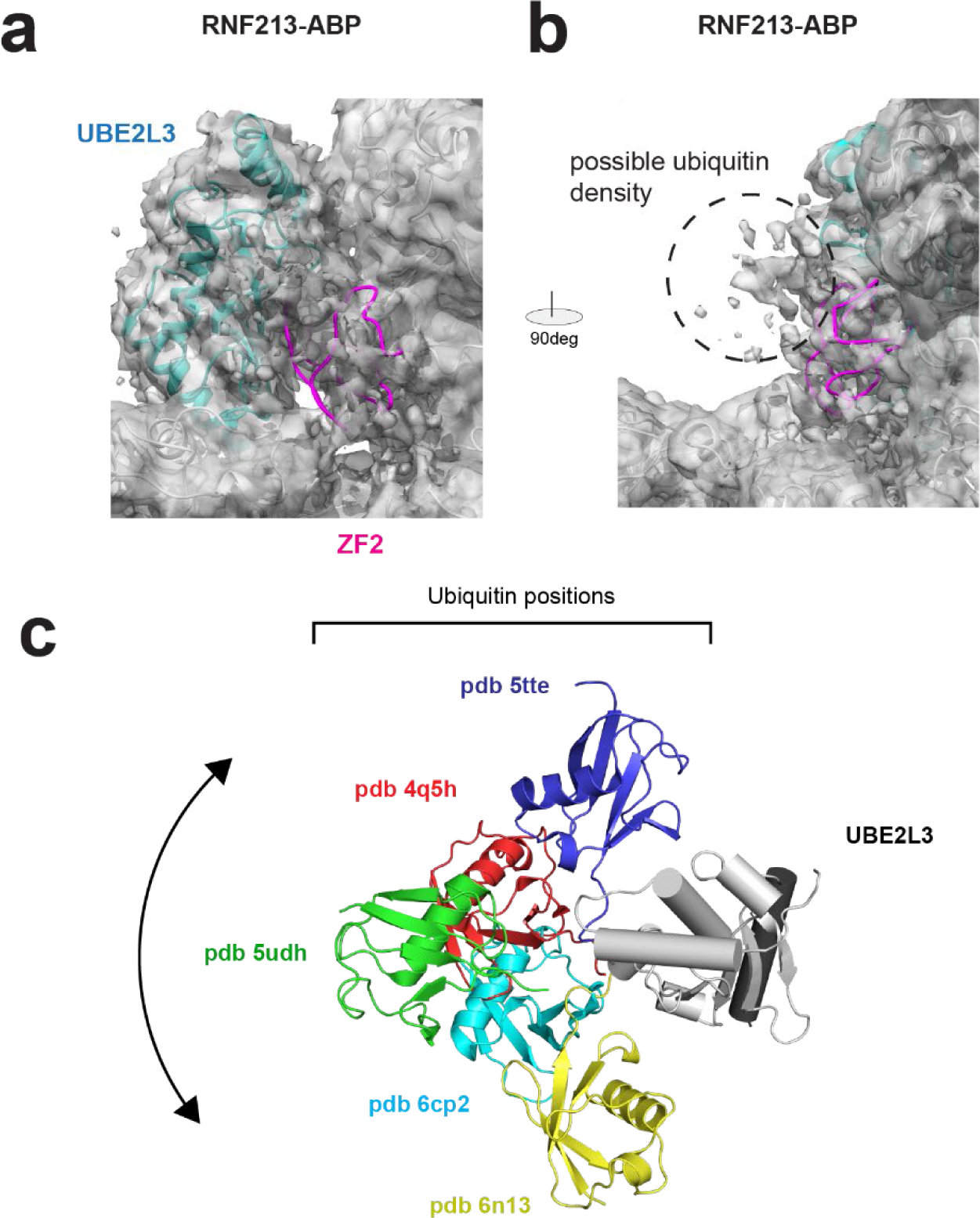
(a) Fitting of the ZF2 domain into the low-resolution density between the main RNF213 body and UBE2L3 in the reconstructed RNF213-ABP cryo-EM map. (b) Low-contour-level density on the side of the UBE2L3, fitting to a possible location of the flexibly attached ubiquitin. (c) Position of ubiquitin (in color) relative to UBE2L3 (light grey) in known structures. Ubiquitin does not have a unique defined relative to UBE2L3, indicating it is likely that the Ub moiety on RNF213 is also flexible. Helix H1 of UBE2L3 is highlighted in dark grey.

**Figure 6-figure supplement 1.**
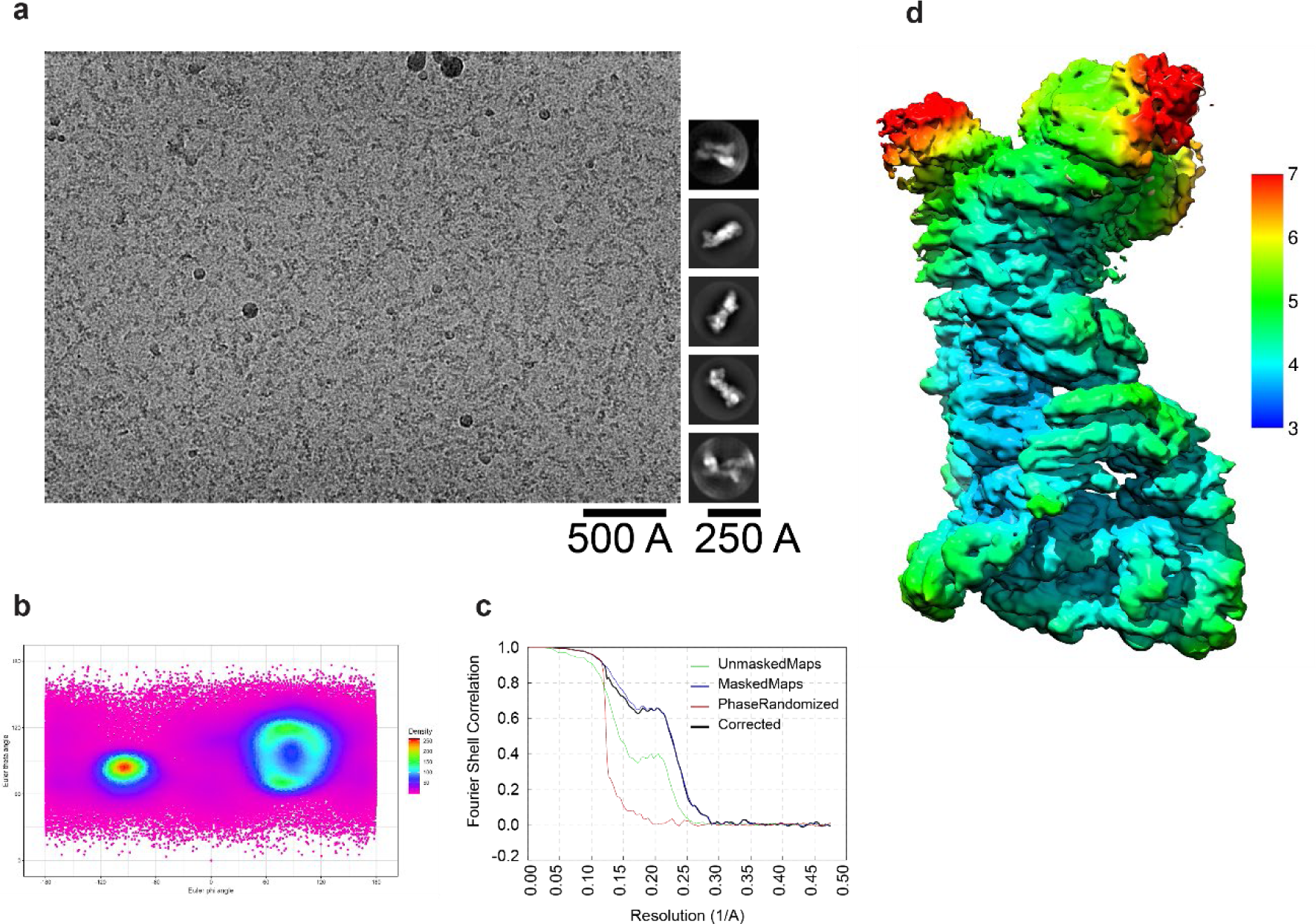
Cryo-EM analysis of RNF213-ABP. (a) Left: Example micrograph, low-pass filtered at 20 Å for visual clarity. Right: Image class averages as generated by 2D classification in relion. (b) Angular distribution heat map of particles used to reconstruct the cryo-EM density. Two clusters of calculated angles are clearly distinguishable, from tilted or non-tilted datasets. (c) Fourier Shell Correlation (FSC) curves indicating a resolution of 4.0 Å using the FSC=0.143 criterion. (d) Reconstructed cryo-EM density colored by local resolution. The resolution color scale is from 3 Å (blue) to 7 Å (red), matching the scale in Figure 5-figure supplement 1.

**Figure 6-figure supplement 2.**
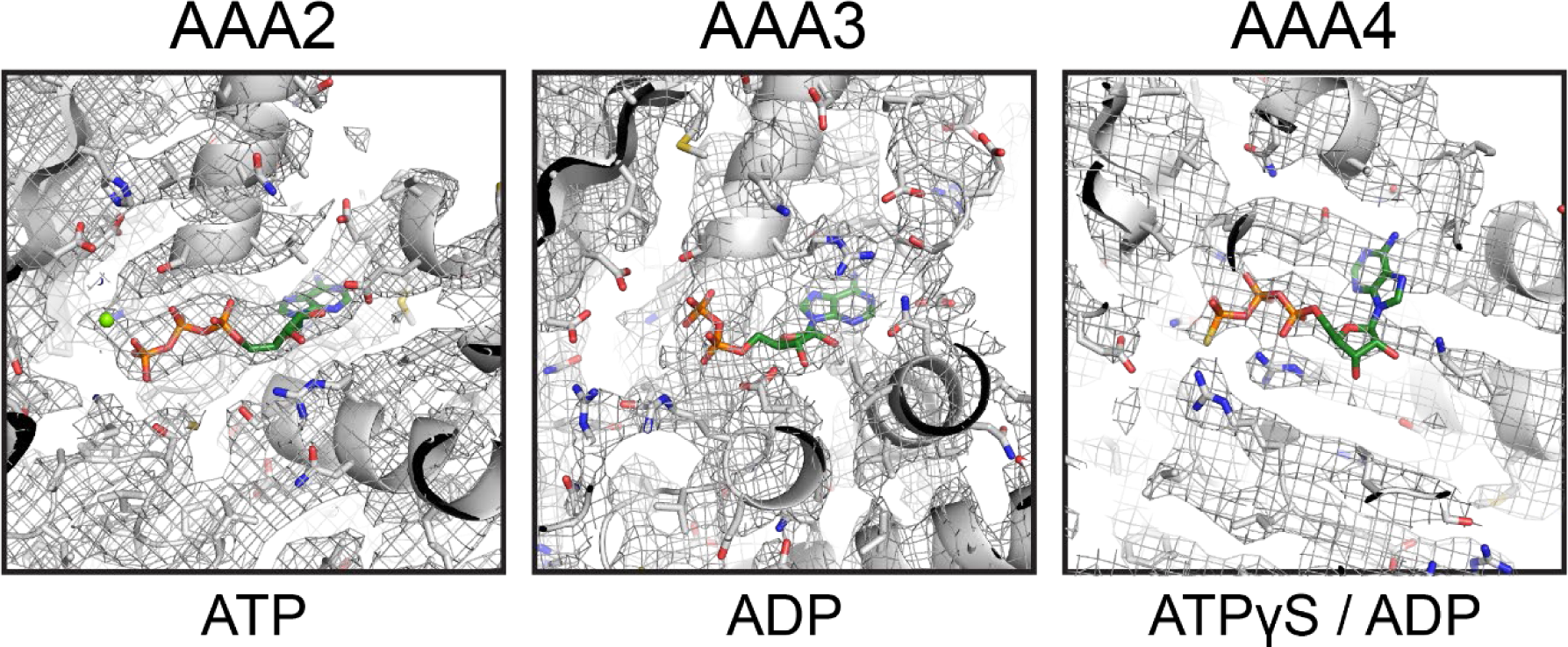
ATP binding sites of the three AAA domains that contain a Walker A motif. Experimental density present in the pocket is shown in orange. In ATPase subunit AAA2, an ATP or ATPγS molecule can be fit unambiguously. AAA3 site density corresponds to an ADP molecule. AAA4 site density corresponds to ATPγS, but with a lower occupancy of the γ-thiophosphate, indicating partial substitution with ADP.

